# Multi-omics profiling of living human pancreatic islet donors reveals heterogeneous beta cell trajectories toward type 2 diabetes

**DOI:** 10.1101/2020.12.05.412338

**Authors:** Leonore Wigger, Marko Barovic, Andreas-David Brunner, Flavia Marzetta, Eyke Schöniger, Florence Mehl, Nicole Kipke, Daniela Friedland, Frederic Burdet, Camille Kessler, Mathias Lesche, Bernard Thorens, Ezio Bonifacio, Cristina Legido Quigley, Philippe Delerive, Andreas Dahl, Kai Simons, Daniela Aust, Jürgen Weitz, Marius Distler, Anke M Schulte, Matthias Mann, Mark Ibberson, Michele Solimena

## Abstract

Existing studies do not sufficiently describe the molecular changes of pancreatic islet beta cells leading to their deficient insulin secretion in type 2 diabetes (T2D). Here we address this deficiency with a comprehensive multi-omics analysis of metabolically profiled pancreatectomized living human donors stratified along the glycemic continuum from normoglycemia to T2D. Islet pools isolated from surgical samples by laser-capture microdissection had remarkably heterogeneous transcriptomic and proteomic profiles in diabetics, but not in non-diabetic controls. Transcriptomics analysis of this unique cohort revealed islet genes already dysregulated in prediabetic individuals with impaired glucose tolerance. Our findings demonstrate a progressive but disharmonic remodeling of mature beta cells, challenging current hypotheses of linear trajectories toward precursor or trans-differentiation stages in T2D. Further, integration of islet transcriptomics and pre-operative blood plasma lipidomics data enabled us to define the relative importance of gene co-expression modules and lipids positively or negatively associated with HbA1c levels, pointing to potential prognostic markers.

## Introduction

Type 2 diabetes (T2D) mellitus collectively defines a cluster of genetically complex pathological states characterized by persistent hyperglycemia, often leading to cardiovascular complications, kidney failure, retinopathy and neuropathies. Affecting more than 450 million people, with rising incidence rates over the past decades, this syndrome is a major threat for public health and society globally^1^. Common determinant and ultimate cause of T2D is the inability of pancreatic islet beta cells to secrete insulin in adequate amounts relative to insulin sensitivity, in the absence of evidence for their autoimmune destruction or a monogenetic deficit. Beta cell failure typically results from a lengthy process spanning many years. Remarkably, however, it can be rapidly reverted upon bariatric surgery or severe caloric restriction^2,3^. These observations argue against the occurrence of major beta cell apoptosis in T2D, especially since adult beta cells hardly replicate, while robust evidence of beta cell neogenesis after puberty is also lacking. Hence, the prevailing opinion is that persistent metabolic stress drives mature beta cells to phenotypically de-differentiate into progenitor cells or trans-differentiate into other islet endocrine cell types over time^4-6^. As the pathogenesis of beta cell dysfunction in T2D remains largely unclear, the diagnosis of this disease relies on accepted, but still surrogate parameters and cutoffs that have been primarily developed for clinical practice to optimize therapeutic interventions^7^.

Insight into molecular alterations associated with impaired insulin secretion in T2D has been largely obtained from pancreatic islets isolated enzymatically from brain-dead or cadaveric subjects classified according to a categorical division into non-diabetic and diabetic, rather than on a continuum from euglycemia to steady hyperglycemia. This approach has multiple shortcomings^8^. Briefly, islet researchers do not generally have access to extensive clinical and laboratory information about the donors prior to their admission to an intensive therapy unit^9^. Moreover, the islet state is perturbed by the metabolic stress associated with a terminal condition and the related pharmacological treatments^10,11^. Enzymatic isolation of islets and their in vitro culture can further change their molecular profile^12,13^. In the attempt to overcome, at least in part, these limitations, we established a complementary platform for the procurement of islets which relies on the collection and analysis of pancreatic specimens from metabolically profiled living donors undergoing pancreatectomy for a variety of disorders^8,14^. We showed that this approach is very reproducible and scalable and provides a novel view on transcriptomic and functional alterations in pancreatic islets of subjects with T2D^15-17^.

The aim of the present study has been to profile in greater detail gene expression changes occurring along the progression from euglycemia to long-standing T2D in human islets *in situ* and to integrate this knowledge with clinical traits, circulating lipid levels and the islet proteome, hence enabling inferences about the mechanisms driving islet dysfunction and the identification of potential biomarkers for it.

## Results

### Recruitment of a large cohort of living donors for islet and plasma omics data

To gain insight into the history of islet cell deterioration along the progression from normal glycemic regulation to T2D, we collected surgical pancreatic tissue samples from 133 metabolically phenotyped pancreatectomized patients (PPP). Eighteen were non-diabetic (ND), 41 had impaired glucose tolerance (IGT), 35 Type 3c Diabetes (T3cD) and 39 T2D (Fig. 1A and Fig. 1B). These group assignments were based on glycemic values at fasting and at the 2 h time point of an oral glucose tolerance test (OGTT) using the thresholds defined in the guidelines of the American Diabetes Association^7^, or, when applicable, on a previously established diagnosis of T2D. In this cohort, 51.9% were males and the mean age was 65.36±11.54 years, with ND PPP being on average younger than the other three groups (Fig. 1C and Supplementary Table S1). The body mass index (BMI) was significantly lower in ND compared to IGT, T3cD and T2D PPP. The HbA1c value, as a parameter of longer-term glycemia, was 5.25±0.3 in ND, 5.75±0.42 in IGT, 6.29±0.95 in T3cD and 7.41±1.29 in T2D PPP (Fig. 1C and Supplementary Table S1). Moreover, based on histopathology, malignant tumors occurred in 50%, 60.97%, 74.29% and 69.23% of ND, IGT, T3cD, and T2D PPP, respectively (Supplementary Table S1).

**Figure 1:**
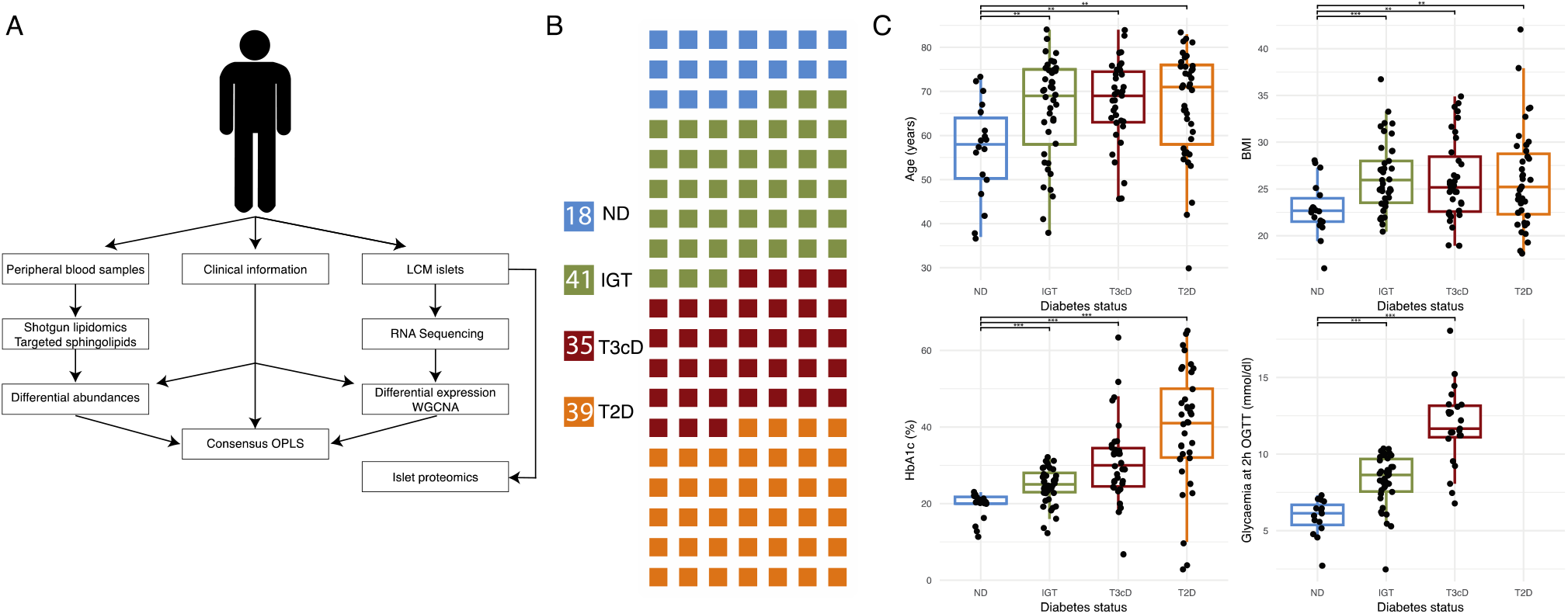
Overview of the experimental procedures and cohort characteristics. A) Experimental procedures overview. Clinical data and peripheral blood were collected preoperatively, and the snap-frozen surgical pancreatic tissue used for LCM of the islets of Langerhans. Blood samples were analyzed for lipidomics, while LCM islets for transcriptomics and proteomics. Omics datasets were individually evaluated in relationship to glycemic status and further integrated with each other using Consensus Orthogonal Partial Least Squares (OPLS) analysis. B) Waffle plot showing the structure of the cohort in terms of glycemic/diabetes categories based on American Diabetes Association criteria. Absolute numbers for each category are given in the legend boxes. C) Boxplots of four major clinical parameters relevant for diabetes diagnosis and management. Statistically significant differences from ND PPP were determined using the Student’s t-test (\**p*<0.05; \*\**p*<0.01). LCM Laser Capture Microdissection, ND Non-diabetic, IGT Impaired Glucose Tolerance, T3cD Type 3c Diabetes, T2D Type 2 Diabetes.

### Pancreatic islet gene expression and pathways drift progressively with glycemia deterioration

Gene expression profiles of islets isolated by laser capture microdissection (LCM) from resected and snap-frozen pancreas samples of ND, IGT, T3cD and T2D PPP were assessed by RNA sequencing. After removal of genes with low expression levels, the overall islet transcriptome encompassed 19,119 genes, of which 14,699±693 were present (raw read counts >0) in ND PPP, 14,967±455 in IGT PPP, 14,939±493 in T3cD PPP and 14,997±428 in T2D PPP. Genes with a fold change (FC)>1.5 and a false discovery rate (FDR)>;0.05 were considered to be differentially expressed (DE) between the groups. Pairwise group comparisons of IGT vs. ND, T3cD vs. ND and T2D vs. ND revealed an exacerbation of gene dysregulation with deterioration of glycemic control (Fig. 2A). Notably, no DE islet genes were identified between IGT vs. ND, while 161 and 650 DE genes were found between T3cD vs. ND and T2D vs. ND, respectively (Fig. 2A and Supplementary Table S2).

**Figure 2:**
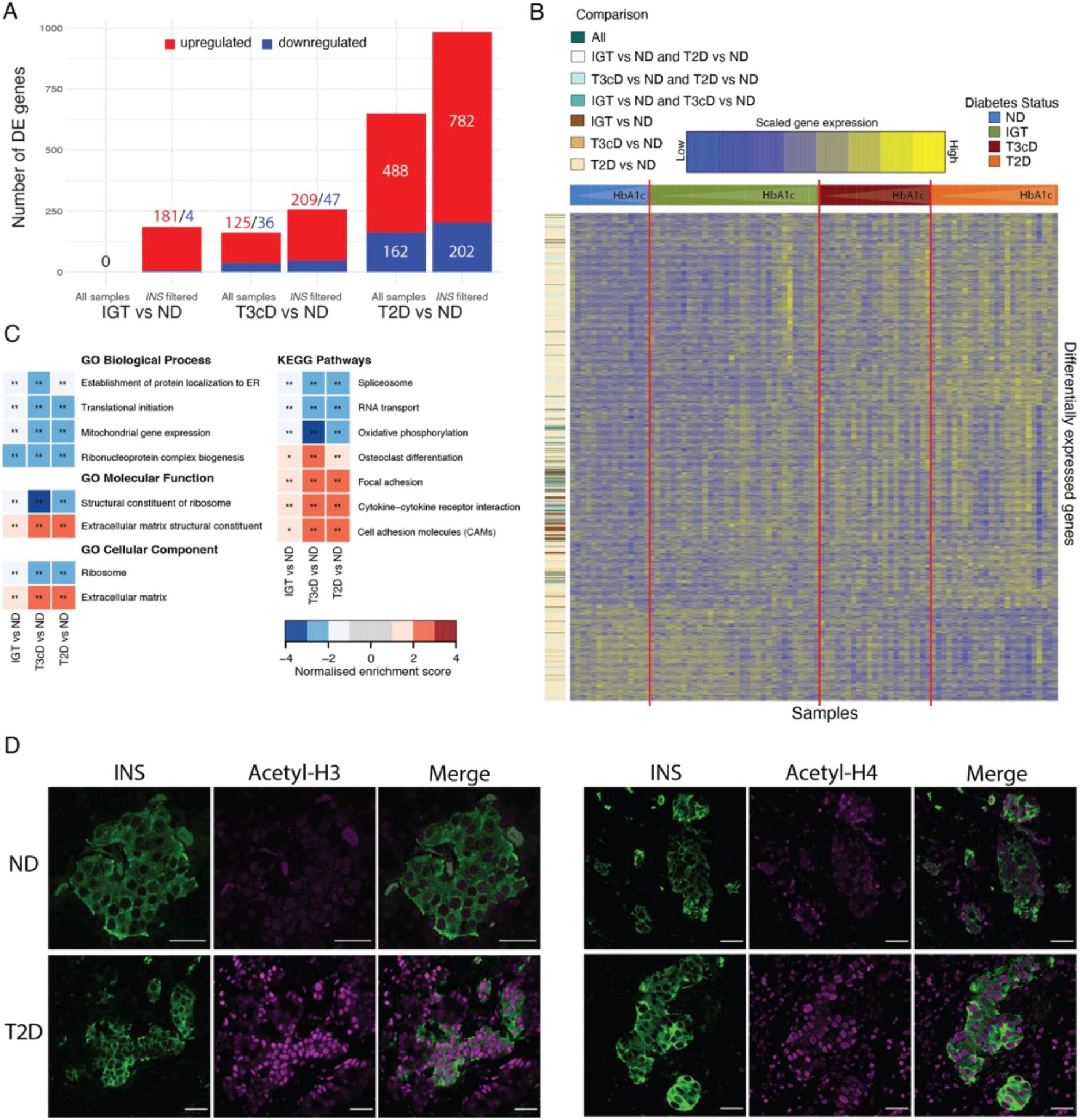
Transcriptional changes between non-diabetic, pre-diabetic and diabetic patients. A) Number of DE genes identified by comparing glycemic groups of PPP in the entire (all samples) or “restricted” cohort (*INS* filtered). B) Gene expression profile of DE genes in the “restricted” cohort. Columns represent patients grouped according to their glycemic status and ordered based on increasing HbA1c levels. Rows, representing DE genes (variance stabilizing transformation normalized counts), were clustered based on Euclidean distance. The colored side bar indicates in which comparisons a gene was identified as differentially expressed. C) Gene Set Enrichment Analysis of DE genes between IGT, T3cD or T2D and ND PPP in the “restricted” cohort. GO terms and KEGG pathways are colored according to the normalized enrichment score. Corresponding p-values are also indicated (\**p*<0.05, \*\**p*<0.01). D) Immunofluorescence for insulin (green), acetylated histones H3 (left) and H4 (right) (magenta) in representative samples of formalin fixed paraffin embedded pancreatic tissues from ND and T2D PPP. Scale bars correspond to 20μm. DE differentially expressed, ND Non-diabetic, IGT Impaired Glucose Tolerance, T3cD Type 3c Diabetes, T2D Type 2 Diabetes.

Restricting the transcriptomic analysis to libraries in which insulin *(INS)* was the most expressed gene resulted in the retention of islet datasets from 15 ND, 35 IGT, 21 T3cD and 24 T2D subjects, without dramatically affecting the overall composition of the cohort in regards to diabetes status and major descriptive parameters (Supplementary Table S3). Deconvolution analysis indicated that in 97.8% of retained samples the proportion of beta cells was >50% (Supplementary Fig. S1), supporting the choice of this strategy to discriminate samples especially enriched in beta cell transcripts. Despite the expected reduction in statistical power due to ~ 30% smaller size of this “restricted” cohort (92 samples retained from 133), the number of DE genes between islets of T2D vs. ND PPP increased by 51% to 984 (782 up, 202 down), and by 59% to 256 (209 up, 47 down) between islets of T3cD vs. ND PPP (Fig. 2A, Supplementary Table S4). Seven of the 984 DE genes are known risk genes for T2D, two upregulated (*SGSM2* and *BCL2*) and five downregulated *(RASGRP1, G6PC2, SLC2A2, ZMAT4* and *PLUT*)^18^, while most of the remaining genes have not been previously reported to be altered in islets of subjects with _T2D_^14,19^.

Among the DE genes in islets of T2D PPP, *INF2* and *AKR7L* were negatively correlated in a moderate fashion with duration of the disease measured in years (Spearman correlation coefficient −0.32 and −0.41 respectively), albeit they were both upregulated relative to islets of ND PPP. Most notably, this filtering step enabled, for the first time, the identification of 185 DE genes between islets of IGT vs. ND PPP. Most of these DE genes were upregulated (181/185), and 98 also dysregulated with the same directionality (97 up, 1 down) between islets of T2D vs. ND PPP. Intriguingly, and apparently at variance with previous eQTL findings^20^, the T2D risk gene *ARAP1* and its neighboring gene *STARD10* were both upregulated and among the 77 genes dysregulated in islets of IGT PPP only. No islet cell type specific genes^21^ were enriched in any of the differential expression analyses. Furthermore, no shift of islet cell type proportions with the progression of the disease was observed in the deconvolution analysis (Supplementary Fig. S1A).

For both the “restricted” and the full data set, heatmaps of gene expression levels in the four patient groups were prepared as a visual complement to the statistical analysis (Fig. 2B and Supplementary Fig. S2A). Despite the marked differences between the findings in the “restricted” and complete cohort, upregulation prevailed as the direction of gene dysregulation in both of them (Fig. 2A and Supplementary Fig. S2A). Based on these observations, pancreatic tissue sections of 5 ND and 5 T2D PPP with the “restricted” cohort were immunostained with antibodies specific for histone H3 and H4 lysine acetylation – an epigenetic modification associated with greater access of transcription factors to promoter sites resulting in increased gene expression. Notably, the immunoreactivity for both acetylated histones was remarkably increased in the islets, and also in the surrounding exocrine cells of T2D PPP, and to a lesser extent IGT PPP (not shown), compared to ND PPP (Fig. 2D).

### Extracellular matrix and mitochondrial pathways are perturbed in T2D and IGT subjects

We further analyzed differentially expressed gene functions by gene set enrichment analysis using Gene Ontology terms and KEGG pathways (Fig. 2C, Supplementary Fig. S2B and Supplementary Tables S5 and S6). Results obtained from the different gene set collections cross-validated each other, since similar biological themes emerged. Islets of pre-diabetic and diabetic subjects displayed upregulation of islet genes that were functionally related to cell-extracellular matrix interaction, immune response and signaling pathways, while expression of genes related to RNA processing, protein translation and mitochondrial oxidative phosphorylation were downregulated. Importantly, the analysis performed on the “restricted” cohort, differently from the full dataset, also revealed that the strength of the enrichment increased with progression of the disease (Fig. 2C and Supplementary Fig. S2B). These data suggest that early dysregulation of gene pathways exacerbates with the decline of beta cell function.

### Weighted gene co-expression network analysis identifies islet gene modules correlated with the elevation of HbA1c

To globally interpret transcriptomic data and identify sets of genes likely to be functionally related and co-regulated, we grouped genes based on similarities in their expression profiles into modules using a network-based approach^22^. In the cohort of 133 PPP, we identified 36 co-expressed gene modules, which were arbitrarily labeled M1 through M36. The expression profiles of the genes in each module were summarized by a module eigengene, or first principal component of the expression matrix. Module eigengenes were used to computationally relate modules to one another and to genes or clinical variables. Correlation between module eigengenes and diabetes-related clinical traits revealed modules M9 and M14 as those with the highest positive and negative correlation with HbA1c, respectively (Fig. 3A and Supplementary Table S7). The former consisted of a set of genes that showed similar patterns of increased expression in most PPP with T2D (Fig. 3B), while the latter was mostly composed of genes with coordinated down-regulation in diseased subject samples (Fig. 3C).

**Figure 3:**
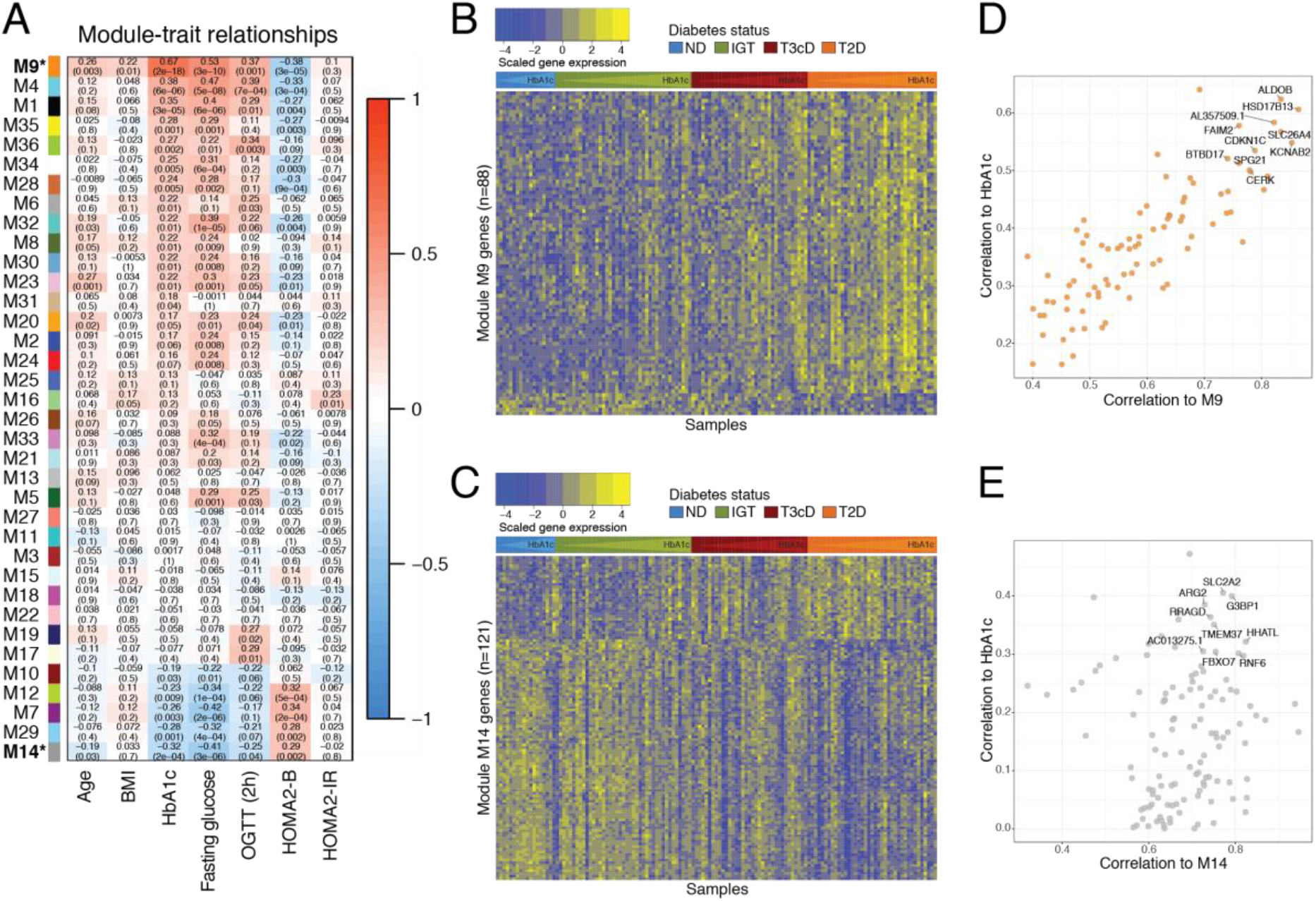
Identification of co-expressed gene modules related to diabetes traits. A) Correlation between module eigengenes and clinical traits including age, BMI, HbA1c, fasting glucose, OGTT at 2 hours, HOMA2-B and HOMA2-IR. Each cell contains the corresponding Spearman correlation coefficient and Student *p* value (in parenthesis). Cells are colored according to their correlation to clinical traits. Modules are ordered based on their correlation to HbA1c. B-C) Gene expression profiles of gene modules M9 (B) and M14 (C). Columns, representing PPP, were grouped according to their glycemic status and ordered based on increasing HbA1c levels. Rows, representing genes (variance stabilizing transformation normalized counts), were clustered based on Euclidean distance. D-E) Scatter plot of module membership vs. gene significance for HbA1c in modules M9 and M14. Genes with the highest module membership and gene significance (“hub genes”) are labeled. ND Non-diabetic, IGT Impaired Glucose Tolerance, T3cD Type 3c Diabetes, T2D Type 2 Diabetes.

We next evaluated how close a gene was to a given module, denoted as module membership, by correlating its expression profile with the module eigengene. This analysis allowed us to identify highly connected genes or “hub” genes for HbA1c-related modules (Fig. 3D-E). These included genes that we had previously identified as differentially expressed in subjects with T2D^14,15^, and which were correlated with HbA1c either positively, such as module M9 genes *ALDOB* (FC=8.45 with adj. *p*<0.001 in T2D vs. ND in “restricted” cohort) and *FAIM2* (FC=7.11 with adj. *p*<0.001 in T2D vs. ND in “restricted” cohort) or negatively, such as module M14 genes *SLC2A2* (FC=-2.77 with adj. *p*<0.001 in T2D vs. ND in “restricted” cohort) and *TMEM37* (FC=-1.73 with adj. *p*<0.001 in T2D vs. ND in “restricted” cohort). Interestingly, we (Supplementary Fig. S3A) and others^23^ found *ALDOB* to be upregulated in islets from 13-week-old diabetic *db/db* mice compared to the heterozygous *db/+* littermate (Supplementary Fig. S3A) as well as in a mouse beta, but not alpha, cell line upon exposure to high glucose (Supplementary Fig. S3B). However, the overexpression of *ALDOB* in beta cells of T2D PPP could neither be verified by in situ hybridization using the RNAScope platform (data not shown), nor by immunofluorescence on tissue sections due to the cross-reactivity of the available anti-ALDOB antibody with other aldolase isoforms (Supplementary Fig. S3C).

### Proteomics of LCM-isolated pancreatic islets reveals heterogenous profiles of T2D subjects and extends target identification

To verify and extend the transcriptomic data at the functional level of proteins, we analyzed the mass spectrometry (MS)-based proteomic profiles of LCM pancreatic islets from five ND and five T2D PPP (Supplementary Table S8). Using a very high sensitivity workflow on a novel trapped-ion mobility Time of Flight mass spectrometer^24^, we identified 2,237±499 islet proteins for ND PPP and 1,819±412 islet proteins for T2D PPP (Figure 4A). Quantitative reproducibility between biological replicates was high with Pearson correlations ranging from 0.83 to 0.95 (Supplementary Fig. S4A). Principal component analysis (PCA) clustered the data into two distinct groups matching the clinical stratification (Fig. 4B). Interestingly, islets of ND PPP clustered closely, indicating a very similar proteome signature, while those of T2D PPP revealed substantial proteome heterogeneity among each other. Differential expression analysis confirmed that islets of T2D and ND PPP have very different proteomic profiles. The main differential drivers are well-characterized markers of pancreatic islet cells, including SLC2A2^25^, and many proteins implicated in mitochondrial structure, translation, energy supply and amino acid or fatty metabolism such as YMEL1, MRPL12, BA3(C14orf159), ACADS and its paralogue ACADSB, which were highly depleted in islets of T2D PPP (Fig. 4C). Besides AKR7L, ACADS was the only other upregulated and differentially expressed gene in islets of both IGT and T2D PPP, while being also downregulated at the protein level. All differentially expressed mitochondrial proteins are encoded by the nuclear genome (Fig. S4B). Intriguingly, the level of the sulfonylurea receptor ABCC8 subunit^26^ was also strongly reduced in islets of T2D PPP. This downregulation might be an effect secondary to pharmacological treatment, as three among these patients had been treated with anti-diabetic SUR1 antagonists glibenclamide (DP197), glimepiride (DP118) or mitiglinide (DP087) (Supplementary Fig. S4C). We found the glycolytic enzyme ALDOB to be on average four-fold upregulated in islets of T2D vs. ND PPP. This is consistent with our transcriptomic data (ALDOB FPKM: 76.16±50.82 in T2D PPP vs. 4.63±0.95 in ND PPP; p=0.03) and with previous^14,15^ and our current WGCNA analyses. Other proteins robustly overexpressed in islets of T2D PPP included the alpha-L-fucosidase FUCA1 and the surface marker for hematopoietic stem cells THY1.

**Figure 4:**
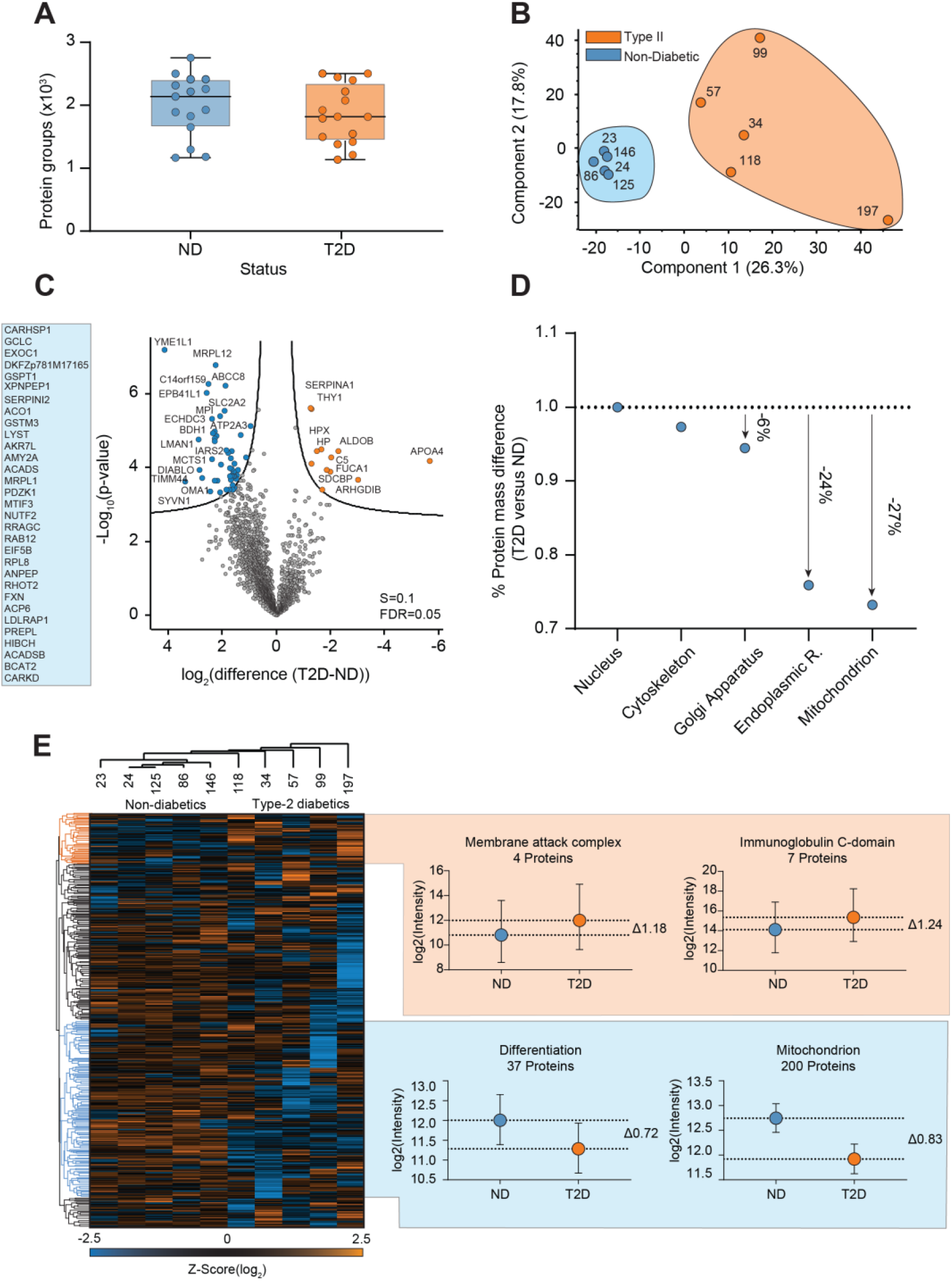
Proteomics Analysis. A) Number of identified proteins from pooled human pancreatic islet cells isolated by LCM from PPP classified as non-diabetic (ND, N=5) or with T2D (N=5). B) Principal Component Analysis (PCA) of all grouped pancreatic islet measurements (ND=blue, T2D=orange). C) Volcano plot comparing *p* values and log2-fold changes between islets of ND and T2D PPP. D) Percentage distribution of total protein islet mass and its contribution per organelle between ND and T2D PPP. The ND/T2D islet protein mass ratio in different organelles was normalized by the nucleus protein mass. E) Hierarchical clustering of all islet proteins identified in the T2D and ND PPP clusters. Log2-transformed intensity values were normalized by z-scoring before the clustering followed by one-dimensional gene ontology enrichment for cellular compartment and keywords for each of the clusters. ND Non-diabetic, T2D Type 2 Diabetes.

Next, we employed the proteomic ruler algorithm and annotations of subcellular localization to compare the protein mass distribution of major cellular compartments^27^ (Fig. 4D). Islets of T2D PPP lost an estimated protein mass of 6% in the Golgi apparatus, 24% in the endoplasmic reticulum, and 27% in the mitochondria compared to those of ND PPP, while cytoskeleton protein mass was unchanged. Unsupervised hierarchical clustering of all 2,622 detected proteins, clustered the data according to clinical categories (Fig. 4E). One-dimensional gene ontology enrichment^28^ revealed two distinct clusters whose protein intensity levels associated with the terms ‘membrane attack complex’ (p<2.18E-04) and ‘Immunoglobulin C-domain’ (p<2.68E-06) were enriched by 2.27-fold and 2.36-fold in islets of T2D vs. ND PPP, respectively. Proteins with the gene ontology-term ‘differentiation’ (p<3.09E-04) and ‘mitochondrion’ (p<2.19E-08) were 1.65 and 1.78-fold in islets of ND PPP.

### T2D patients show decreased levels of plasma phospholipids and elevated levels of plasma (dihydro-)ceramides

Our study encompassed two independently generated lipidomics data sets. First, shotgun lipidomics was performed on peripheral blood plasma samples of the aforementioned cohort (4 ND, 21 IGT and IFG, 13 T3cD and 17 T2D) (Supplementary Tables S9 and S10). Second, sphingolipid profiling was performed on peripheral blood samples of subjects within the cohort subjected to transcriptomic analysis (11 ND, 32 IGT and IFG, 26 T3cD and 32 T2D) (Supplementary Tables S11 and S12). Prior to data analysis, lipidomics samples from PPP with very high bilirubin values (>100 μmol/l) were removed to avoid bias in lipidomics profiles. All available samples from non-diabetic PPP (ND, as previously defined) and the subset of IGT PPP with HbA1c<6.0 were combined into one group, which is referred to here as ND for readability.

In shotgun lipidomics, 113 lipid species from 11 classes were included in the data analysis. When comparing T2D and T3cD to ND PPP, the majority of lipid classes displayed a remarkably homogeneous downward-trend of the individual lipid species they comprised (Fig 5A-B). Most prominently, plasma concentrations of lipids within the phosphatidylcholine (PC O-) class, a large class with 30 measured species, were lower in T2D versus ND PPP. Sixteen lipids of this class were significantly decreased (adjusted p<0.05) after adjusting for age and sex differences, with all of them showing at least a 1.4-fold change. Two lipid species from two smaller phospholipid classes (lysophosphatidylcholines (LPC) and phosphatidylinositols (PI)), and one from the sphingomyelin class (SM), were also significantly less abundant in T2D than in ND PPP (LPC 18:0;0: FC=-1.54, adj. p=0.03; PI 18:0;0/18:2;0: FC=-1.36, adj. p=0.04; SM 34:1;2:, FC=-1.24, adj. p=0.04). (Fig. 5A-B and Supplementary Table S13).

**Figure 5:**
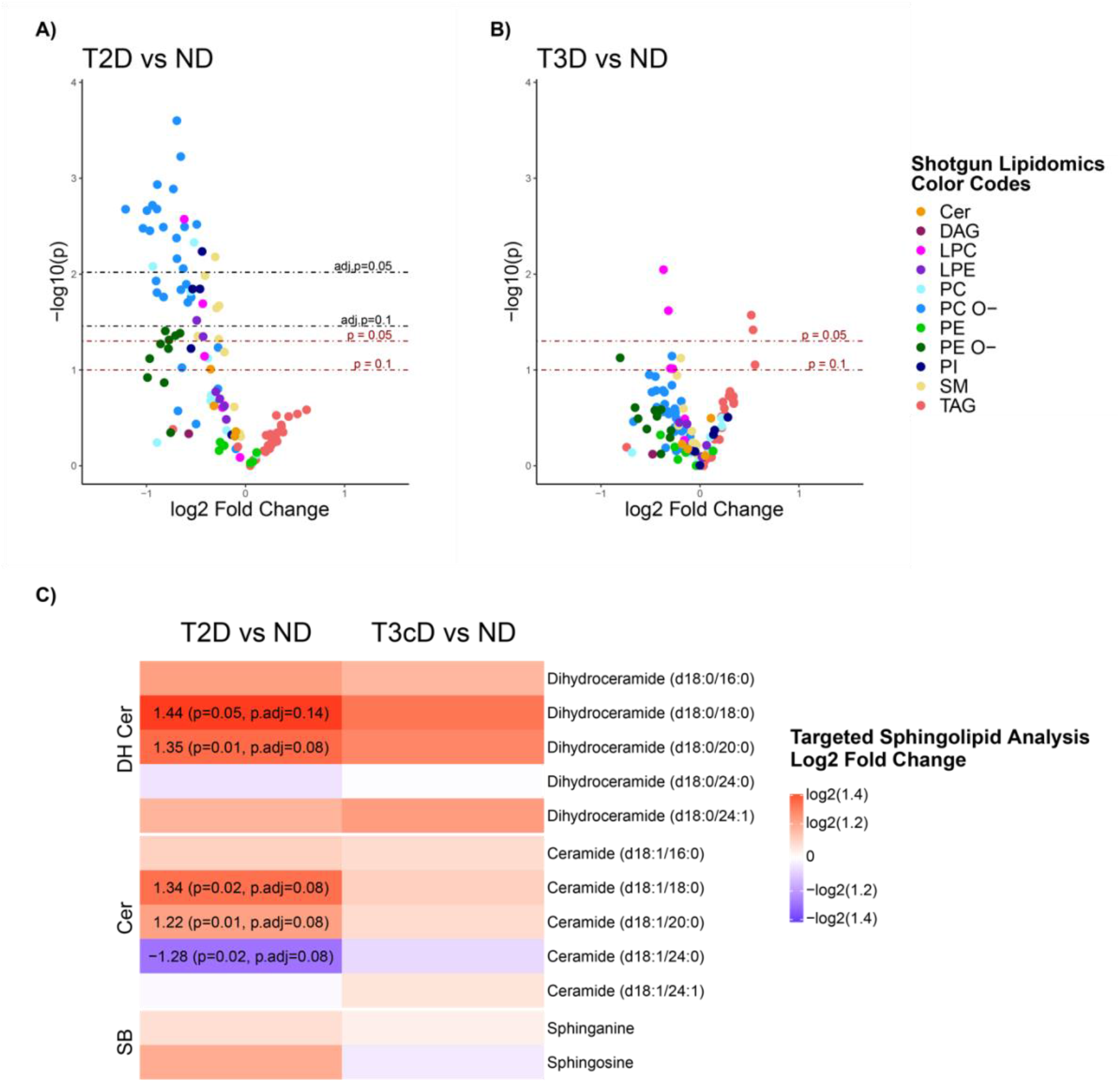
Results from lipidomics differential analysis. A-B) Shotgun lipidomics covering a variety of lipid classes: Ceramides (Cer), Diacylglycerols (DAG), Lysophosphatidylcholines (LPC), Lysophosphatidylethanolamines (LPE), Phosphatidylcholines (PC), Ether-linked Phosphatidylcholines (PC O-), Phosphatidylethanolamines (PE), Ether-linked Phosphatidylethanolamines (PE O-), Phosphatidylinositols (PI), Sphingomyelins (SM), Triacylglyerols (TAG). Volcano plots represent comparisons of plasma lipid levels between ND and T2D PPP. The X-axis shows direction and magnitude of the change; the Y-axis represents the statistical significance of the change. Each point is a lipid species, colored by lipid class to highlight class-specific trends. C) Targeted lipidomics on dihydroceramides (DH Cer), ceramides (Cer) and Sphingoid bases (SB). Each heatmap column represents the comparisons of plasma levels between ND and T2D PPP. Heatmap colors represent direction and magnitude of the change. Log_2_ Fold Change: ratio of mean lipid concentration in the two groups, log2 transformed. Statistical model used for all panels: linear regression with age and sex as covariates (p: *p* value); adjustment of *p* values across all lipid species by the Benjamini-Hochberg method (adj. *p*: adjusted *p* value). T2D Type 2 Diabetes, T3cD Type 3 Diabetes, ND & PD non-diabetic and pre-diabetic (with impaired fasting glucose and/or impaired glucose tolerance) with HbA1c<6.0.

Next, we performed targeted sphingolipidomics on 14 distinct lipid species for very accurate plasma level estimation (ceramides, dihydroceramides and sphingoid bases). Plasma levels of ceramides d18:1/18:0 and d18:1/20:0 were increased in T2D compared to ND PPP (Cer d18:1/18:0: FC=1.34, p=0.02; Cer d18:1/20:0: FC=1.22, p=0.01). Of note, a similar trend towards elevation in T2D vs ND was also observed in the two dihydroceramide species having the same chain lengths as these ceramides (DH Cer d18:0/18:0: FC=1.44, p=0.05; DH Cer d18:0/20:0: FC=1.35, p=0.01). Thus, in our data set, plasma concentrations of ceramides and their precursor dihydroceramides appear to increase simultaneously in T2D.

### Integrative data modelling identifies cell-matrix interaction, cell signaling and immune response as key pathways linked to pancreatic islet dysfunction

To identify a multivariate molecular profile that explains diabetes progression in the PPP cohort, we performed a large-scale integrative multi-omics analysis combining clinical data with islet transcriptomics and plasma lipidomics. Integration of transcriptomics and lipidomics data in the same model enables to weigh the relative importance of lipid and gene expression features in relationship to a chosen clinical trait. Hence, we explored the relationship between gene co-expression modules and plasma lipids by computing a consensus orthogonal partial least square (consensus OPLS)^29,30^ model with HbA1c as the outcome. All three types of biological data, namely gene co-expression modules, lipids from shotgun analysis and sphingolipids from targeted analysis, contributed to the model (35%, 46.5% and 18.5%, respectively), suggesting that they help to explain HbA1c levels in a complementary way. Among them, different lipids and gene modules appear as the most relevant variables in the statistical modelling of HbA1c levels (Fig. 6A, 6B and Supplementary Table S14). Importantly, the model explained a large portion of data variance, highlighting a good fit with the experimental data (see Methods for more details). Among all considered biological data, the co-expression modules M1, M4, M8, M9, M30, M35 and M36 were the top predictive variables for high HbA1c levels, along with the two ceramide species C20 and C18. TAGs were also contributing, although to a lesser extent (Fig 6A, right hand side). Conversely, low levels of HbA1c were strongly related to the co-expression modules M12 and M14 (Fig 6A, left hand side). However, the majority of the predominant predictors for low HbA1c were lipid species, most importantly the PC O-class. This class was also found to be lower in T2D compared to ND patient groups in differential abundance analysis, as shown in Fig 5A. A number of SM, PI and PC lipid species were next in the importance ranking related to low HbA1c, followed by the gene co-expression module M29. These results suggest that the profile of patients with increased HbA1c is characterized by multiple molecular components, some of which represent signals that were not captured by differential abundance analyses comparing diabetes status groups nor by correlating gene co-expression modules individually to HbA1c. Most importantly, consensus OPLS multi-omics analysis pointed towards additional gene co-expression modules that may play a role in glucose dysregulation.

**Figure 6:**
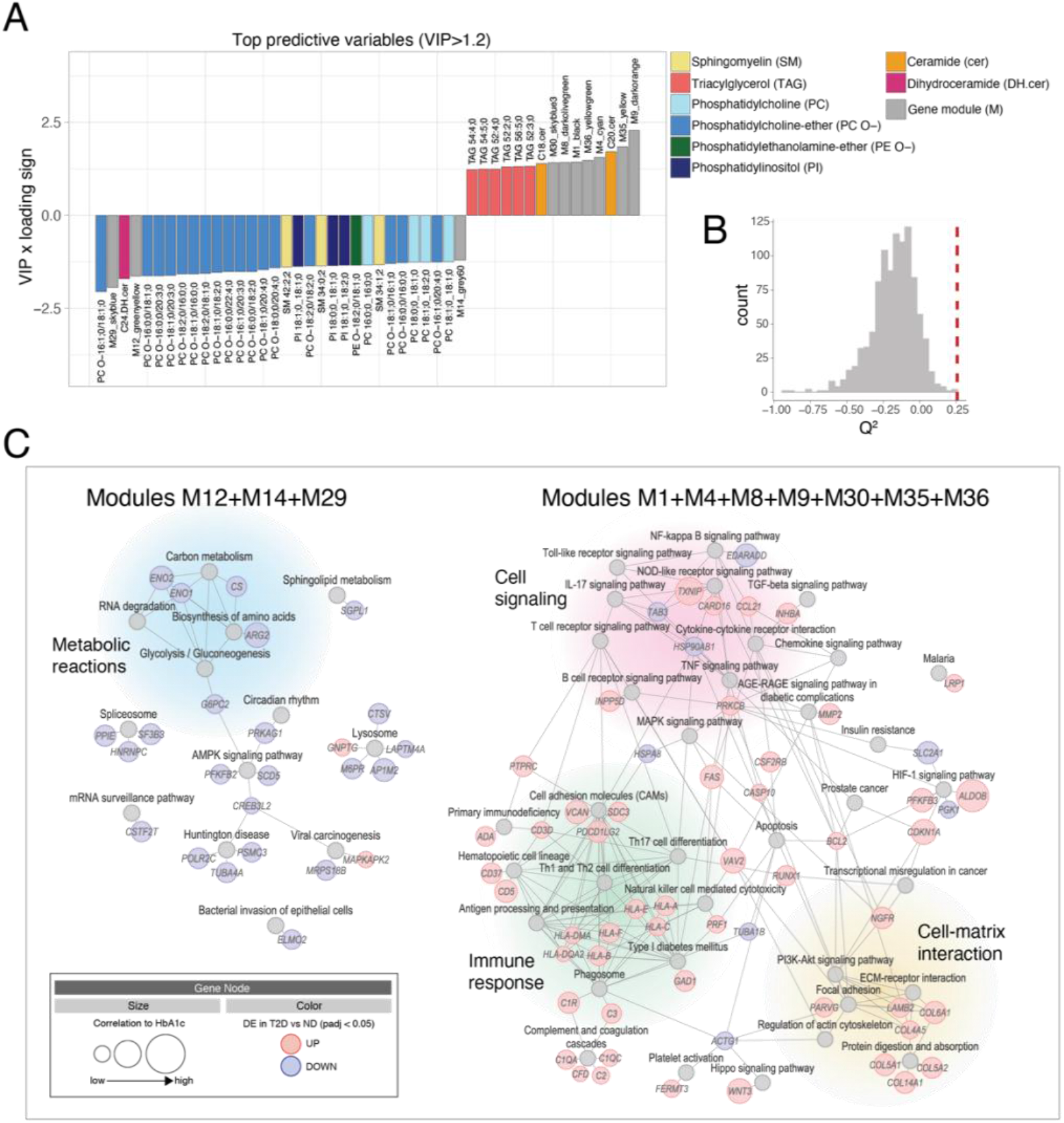
Multiblock data modeling of HbA1c. A) Barplot showing the variable importance in the multiblock consensus OPLS model. The Y-axis represents the importance scores for the predictors multiplied by the sign of the loadings on the predictive latent variable. Variables with importance in projection > 1.2 were selected. B) Statistical significance of the model through permutation test. C) Network representation of functional pathways enriched in modules with best prediction scores for HbA1c. Pathways are represented as gray nodes. Genes are represented as nodes sized based on their correlation to HbA1c and colored based on their differential expression in T2D vs. ND PPP. Only genes with significant differential expression (adj. *p*<0.05) in the “restricted” cohort are shown. VIP Variable Importance in Projection, DE Differentially expressed, ND Non-diabetic, T2D Type 2 Diabetes.

Next, we used the results from the integrative data modelling to infer a network of key altered biological pathways in dysfunctional beta cells. To this end, we pooled gene modules positively associated with HbA1c levels (M1, M4, M8, M9, M30, M35 and M36) (Fig. 6A) and assessed their overlap to KEGG pathways by over-representation analysis. We found that the biological themes underlying these genes were very similar to the pathways upregulated in T2D and IGT PPP and include cell-matrix interaction, cell signaling and immune response (Fig. 6C and Supplementary Table S15). The same strategy was used to identify pathways associated with genes from modules with a negative prediction score for HbA1c (M12, M14 and M29) (Fig. 6A), revealing an enrichment for metabolic pathways (Fig. 6C and Supplementary Table S15).

Of note, several islet genes dysregulated in T2D PPP were driving the enrichment of these pathways. These include, for example, *ALDOB,* which stood out for its strong correlation to HbA1c levels (Fig. 3D and Fig. 6C). These genes, or the proteins encoded by them, should be regarded as putative candidate biomarkers for monitoring disease progression and therapeutic intervention.

## Discussion

This study provides the most extensive dataset on islets *in situ* and plasma samples from the largest cohort of in-depth metabolically profiled living donors. Multi-omics data were generated using state-of-the-art approaches and integrated in a fashion not previously used in studies on islet dysregulation in relation to hyperglycemia in humans. Our transcriptomic and proteomic data from islets *in situ* of ND subjects represent a valuable reference for future investigations. Furthermore, we could identify for the first time a set of islet genes altered in their expression already in subjects with impaired glucose tolerance. This, in turn, enabled us to acquire an unprecedented cross-sectional overview of the progression of islet gene dysregulation in parallel with the continuous elevation of HbA1c values, beyond conventional thresholds for clinical classification of patients.

Pathways involved in RNA biology and especially in mitochondrial function emerged to be most negatively perturbed - a conclusion which in the case of the latter was strongly corroborated by the proteomic analysis, which enabled the identification of known and unknown differentially expressed proteins in islets of T2D PPP. In this context, we emphasize the downregulation of mitochondrial ACADS and its paralogue ACADSB, which catalyze the beta oxidation of short-chain fatty acids, including sodium butyrate. This finding is intriguing in view of the ability of this metabolite to broadly upregulate gene expression through inhibition of histone deacetylases. Unlike in previous studies on isolated islets from brain-dead organ donors^14,18^, the vast majority of differentially expressed genes in islets of T2D, but also IGT and T3cD PPP were upregulated. Among those genes, *ALDOB* stands out being the one with the strongest correlation with the islet gene module M9, which in turn has the strongest correlation with elevated HbA1c. Since *ALDOB* is a marker of beta cell precursors, its overexpression could be interpreted as a sign that in T2D, mature beta cells revert back to an immature stage of differentiation, or that a compartment equivalent to the lifelong niche of virgin beta cells identified in adult mice^31^ expands as a potential compensatory source of new beta cells. However, no additional disallowed gene of immature beta cells, markers of beta cell precursors or other islet cell types were dysregulated, while key determinants of mature beta cells, such as *PDX1, MAFA, NKX6.1* or *UCN3* were unchanged, at least at the transcriptomic level. Retention of fractions of major islet cell types (alpha, beta and delta) within the islet in T2D, consistent with recent imaging studies in samples from pancreatectomized subjects^17^, was confirmed by deconvolution analysis. Our global unbiased proteomic analysis, which corroborated the upregulation of

ALDOB, further showed that the expression profile of islet cells in T2D PPP is very divergent, opposite to its remarkable homogeneity in islet cells of ND subjects. Hence, the regression of beta cells toward a de-differentiated state following a linear trajectory recapitulating their developmental path to maturation or their transdifferentiation into other islet cell types seems less likely than a disharmonic relaxation of constraints on gene expression. Such processes, although possibly reversible, could perturb the coordinated operation of islet cells, including beta cells. In line with this, Lawlor *et al.* reported no evidence of beta cell dedifferentiation/transdifferentiation and alterations in fractions of islet cells in the context of T2D upon sequencing of single islet cells from a small cohort of ND and T2D organ donors^32^. For the future it would be important to assess whether overexpression of ALDOB occurs indeed in beta cells and if it affects their glycolysis and metabolism, taking into account that its paralogue ALDOA, whose RNA and protein levels were unchanged, remains by far the predominant islet aldolase species. Attention may also be directed toward understanding whether impaired oxidative phosphorylation, as a likely outcome of the massively decreased expression of mitochondrial proteins, and thus energy balance homeostasis, accounts, at least in part, for the observed less restrained gene expression.

Similar to findings in other population-based studies on T2D^33,34^, PC O- and LPC lipids were altered in our cohort of T2D PPP, thus supporting the general implications of our observations. In particular, we found that more than half of the PC O-class lipids (16 out of 30) and two of six LPC lipids were lower in T2D compared to ND PPP. In the present study we also found that several ceramides and dihydroceramides are elevated in T2D vs. ND, and whilst these increases were modest, these findings are consistent with those observed in several other recent studies^35-37^, highlighting the importance of these lipids as potential biomarkers of beta cell function in T2D.

Finally, we use a data fusion method^29,30^ to generate a model of how different molecular features (islet gene co-expression, plasma shotgun lipidomics and targeted sphingolipidomics) contribute to HbA1c levels in a continuum from healthy individuals to those with overt T2D. This model allowed us to measure the *relative* importance of different molecular components in explaining HbA1c variability, providing unique insights into the molecular profiles of individuals as they lose glycemic control towards development of T2D. To our knowledge this is the first time such an approach has been used in this field and we suggest that, by modelling multiple levels of information at the same time in deeply phenotyped populations such as the one presented here, we can gain a holistic view of the system and draw conclusions regarding key pathways, targets and biomarkers in metabolic and other diseases.

## Data availability

RNA Sequencing data was deposited in the GEO database with GEO accession number (to be provided once the deposition process is completed)

The proteomics raw datasets and the MaxQuant output files generated and analyzed throughout this study were deposited at the ProteomeXchange Consortium via the PRIDE partner repository with the dataset identifier PXD022561 (https://www.ebi.ac.uk/pride/archive/).

Lipidomics data will be made publicly available shortly.

## Acknowledgement

We wish to thank Leif Groop, Emma Ahlqvist, Stephan Speier and Triantafyllos Chavakis for discussion; Katja Pfriem for administrative assistance. This project has received funding from the Innovative Medicines Initiative 2 Joint Undertaking under grant agreement No 115881 (RHAPSODY). This Joint Undertaking receives support from the European Union’s Horizon 2020 research and innovation program and EFPIA. This work is further supported by the Swiss State Secretariat for Education, Research and Innovation (SERI) under contract number 16.0097-2. The opinions expressed and arguments employed herein do not necessarily reflect the official views of these funding bodies. Work in the Solimena lab is also supported with funds from the German Ministry of Education and Research to the German Center for Diabetes Research (DZD).

## Author contributions

J.W. and M.D., patient recruitment and surgery, provision of clinical data; E.S., N.K. and D.F., sample collection and processing, data entry; D.A., pathology; M.B., N.K. and E.S., patient database management and selection; A-D.B. and M.M., proteomics; M.L., A.D., RNA sequencing, C.L.Q., P.D., K.S., lipidomics and sphingolipidomics; L.W., M.B., A-D.B., F.Ma., F.Me., F.B. and C.K., analysis and integration of multi-omics data; E.B., autoantibody test; A.S., data in mouse tissue and cell lines; M.B., immunofluorescence stainings and antibody validation; B.T., D.A., J.W., A.S., M.M., M.I. and M.S., conceptual insights and provision of funds; L.W., M.B., A-D.B., F.Ma., F.Me., A.S., M.I., M.M. and M.S., writing of the manuscript. All authors read, revised and approved the final version of the manuscript.

## Material and methods

### Cohort

Our cohort comprised 133 adult surgical patients from the University Hospital Carl Gustav Carus Dresden who after informed consent participated in this study over a period of 5 years. Based on the thresholds set by the American Diabetes Association^7^ (ADA) for fasting glucose, HbA1c and 2-hour glycemia of an oral glucose tolerance test (OGTT) in the days immediately before surgery 18 of these patients were classified as non-diabetic (ND), 41 with impaired glucose tolerance (IGT), including 3 with impaired fasting glucose (IFG) only, 35 with Type 3c Diabetes (T3cD) and 39 with Type 2 Diabetes (T2D). A diagnosis of T3cD was made whenever the occurrence of diabetes was not recognized for longer than 1 year prior to the onset of the symptoms leading to surgery and the subject was negative for the presence of circulating autoantibodies against pancreatic islets, which were assessed as previously described^14^. In all analyses IFG and IGT subjects were merged in one group hereinafter labeled as IGT PPP. Medical and family history and relevant clinical biochemistry data available from the routine medical processing of the patients were retrieved from the hospital database and referring physicians. Patients who underwent neoadjuvant chemotherapy as well as those with endocrine neoplasms of the pancreas were excluded from this study.

### Human pancreatic tissue and peripheral blood processing

Surgical tissue specimens were examined by a certified pathologist immediately after resection as per regular clinical procedures. Fragments of healthy pancreatic tissue from the resection margins were excised, snap frozen in liquid nitrogen and stored at −80°C either natively or embedded in TissueTek OCT compound. Estimated warm and cold ischaemia time was on average 2 hours. Peripheral blood samples were stored at −80°C in aliquots of full blood, plasma and serum.

### Transcriptomics

#### Islet procurement and RNA isolation

Pancreatic tissue was sectioned in a cryostat and mounted on UV pre-treated Zeiss MembraneSlide 1.0 PEN slides. Laser capture microdissection (LCM) was done with a Zeiss Palm MicroBeam system using autofluorescence to identify islets, as previously described^38^. RNA was isolated from approximately 20×6μm3 of islet tissue using the Arcturus PicoPure RNA Isolation Kit. Only preparations with RNA Integrity Number ≥5 were used for RNA sequencing. The entire handling of the tissue samples was done in a strictly RNAse free environment.

#### Library preparation, RNA Sequencing and alignment

Sequencing libraries were prepared from bulk RNA using the Illumina SmartSeq protocol.

Single ended 76bp sequencing was done with an Illumina HiSeq 2500 or Illumina HiSeq 500 at the Next Generation Sequencing Core Facility of the CMCB Dresden, with the target depth of 35 million fragments per library. From FASTQ files, purity-filtered reads were trimmed with Cutadapt to remove adapters and low-quality sequences (v. 1.8)^39^. Reads matching to ribosomal RNA sequences were removed with fastq_screen (v. 0.11.1)^40^. Remaining reads were further filtered for low complexity with reaper (v. 15-065)^41^. Reads were aligned against Homo sapiens GRCh38.92 genome using STAR (v. 2.5.3a)^42^. The number of read counts per gene locus was summarized with htseq-count (v. 0.9.1)^43^ using Homo sapiens GRCh38.92 gene annotation. Quality of the RNA-seq data alignment was assessed using RSeQC (v. 2.3.7)^44^.

#### RNA Sequencing quality control, processing and differential expression analysis

RNA Sequencing datasets were screened for exocrine contamination in an initial quality control (QC) step. Analysis of the absolute number of detected expressed genes, gene body coverage and cumulative gene diversity assessment flagged a number of libraries to be of insufficient quality for downstream analysis. Libraries were filtered for minimal expression by removal of genes with less than 5 mean raw reads. Reads were normalized for library size and transformed for variance stabilizing using tools from the DESeq2 Bioconductor package^45^. Further analysis revealed 41 libraries in which transcripts other than insulin (INS) displayed the highest normalized number of reads. Differential expression analysis across the clinical categories (ND, IGT, T3cD, T2D) was performed using limma function with voom approach from limma Bioconductor package^46,47^ on both the full dataset of 133 libraries which passed the QC analysis as well as on the “restricted” dataset of 92 libraries featuring INS as the highest expressed gene based on the linear model with age, gender and BMI as covariates.

#### Gene set enrichment analysis of differentially expressed genes

Functional enrichment analyses of differentially expressed genes in IGT, T2D or T3cD compared to ND patients were performed by weighted gene set enrichment analysis (GSEA) on unfiltered gene lists ranked by decreasing differential expression test statistics. Gene Ontology (GO) term and Kyoto Encyclopedia of Genes and Genomes (KEGG) pathway collections were restricted to gene sets with a minimum and maximum sizes of 100 and 500, respectively. The enrichment scores were normalized by gene set size and their statistical significance was assessed by permutation test (n=1,000). GO enrichment analyses were carried out using the gseGO function from the R package clusterProfiler (version 3.10.1)^48^. GO terms enriched in at least one comparison were identified using *p* value and normalized enrichment score thresholds < 0.01 and > 2.5, respectively. Redundancy of enriched GO terms was removed using the clusterProfiler simplify function (selecting the most representative term by *p* value) and enrichment maps generated using the emapplot function from the R package enrichplot (version 1.2.0). KEGG pathway enrichment analyses were performed using the clusterProfiler gseKEGG function. Results were filtered based on a *p* value threshold < 0.01 and a normalized enrichment score threshold > 2. To simplify results visualization and interpretation, redundant KEGG pathways were also collapsed into fewer biological themes using the enrichment map visualizations.

### Weighted Gene Correlation Network Analysis

#### Gene Co-expression Network Construction

The gene co-expression network was created following the weighted gene correlation network analysis (WGCNA) protocol as implemented in the WGCNA package in R (version 1.68)^22^, as previously described^14^. WGCNA was performed on batch-corrected, normalized and variance stabilizing transformed expression data from the full cohort of 133 subjects. The co-expression network was constructed by calculating an adjacency matrix using Pearson correlation, pairwise complete observations and unsigned method. The soft-threshold parameter was optimized with the function pickSoftThreshold and the best threshold (α = 7) selected by visual inspection. The adjacency matrix was then computed into a topological overlap matrix (TOM), converted to distances, and clustered by hierarchical clustering using average linkage clustering. Modules were identified by dynamic tree cut using the hybrid method and parameters minClusterSize=20 and deepSplit=2. Similar modules were merged using a module eigengene distance of 0.15 as the threshold.

#### Identification of co-expressed gene modules related to diabetes trait

We correlated the module eigengenes to clinical traits using Spearman correlation (pairwise complete observations) and calculated the corresponding *p* values using the cor and corPvalueStudent functions from the WGCNA package, respectively. Module-trait correlations were represented as heatmap using the labeledHeatmap function from the WGCNA package. The modules displaying the most positive or negative correlation to HbA1c were further analysed. Normalized and variance stabilizing transformed gene counts for selected modules were plotted as heatmap using the heatmap.2 function from the R gplots package (version 3.0.1.2). Rows (representing genes) were scaled and hierarchically clustered by Euclidean distances. Columns, representing patients, were custom ordered as described in the legend of figure 3. Module hub genes, such as highly connected genes within a module that could have a strong influence on a phenotypic trait, were identified as those with the highest correlation with the particular trait and the highest correlation with the module eigengene.

#### Significance of gene co-expression modules

We tested the significance of the co-expression modules by comparing their intramodular connectivity (connectivity between nodes within the same module, as computed by the WGCNA intramodularConnectivity function) to the background as follows. For each selected module of size N, we calculated a Z-score as:

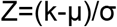

where k is the intramodular connectivity and μ and σ are the mean and standard deviation of the intramodular connectivity from 1,000 randomly sampled modules of size N respectively. Empirical p values were then calculated as the fraction of random intramodular connectivity values ≥ to the observed intramodular connectivity. For the modules with the highest variable importance in projection score in the HbA1c multiblock model, all of the random intramodular connectivity values were below the observed intramodular connectivity, suggesting that these modules were more compact than modules assembled by randomly sampling the same number of genes from the expression data (Supplementary Table S7).

#### Functional profiles of gene modules with best prediction score for HbA1c

The clusterProfiler enrichKEGG function was used to test for the over representation of selected co-expressed gene modules in KEGG pathways using hypergeometric distribution. A *p* value threshold < 0.01 was used to identify enriched terms. Enrichment map visualizations were used to overcome gene set redundancy. Results were displayed as networks of enriched pathways and overlapping genes using cytoscape (version 3.5.1).

#### Deconvolution analysis

In all samples a cell proportions matrix was produced using the R package DeconRNASeq (v.1.26.0) on RPKM-transformed data. The signature file provided to DeconRNASeq comes from Xin et al. (2016)^21^, Supplementary Table S2 (A), obtained using single-cell data. It was adapted to the human genome version 38 by excluding 15 obsolete genes.

### Lipidomics

#### Sample availability and sample overlap with transcriptomics data

Pre-operative plasma lipidomics samples were obtained from a subset of the PPP cohort. Shotgun lipidomics analysis was performed on plasma from 55 PPP. These included 53 subjects who also had their islet transcriptomics profile included in this study plus two PPP who were not part of the transcriptomics analysis because the RNA-Seq data failed to pass the quality control. Moreover, targeted sphingolipid analysis was performed on plasma from 101 PPP. These included 98 PPP whose transcriptomics data was also included in this study plus three PPP whose RNA-Seq data was excluded for quality reasons. The number of samples in the two types of lipidomics analysis was smaller than in islet transcriptomic analysis because of the limited availability of plasma samples.

#### Shotgun lipidomics measurements

A streamlined mass-spectrometry (MS) -based platform for shotgun lipidomics developed by Lipotype GmbH (Dresden, Germany) was used for lipidomic profiling of patient plasma samples. Lipid extraction, internal standard addition and infusion into the mass spectrometer were performed as previously described^49^. The internal standard mixture contained: cholesterol D6 (chol), cholesterol ester 20:0 (CE), ceramide 18:1;2/17:0 (Cer), diacylglycerol 17:0/17:0 (DAG), phosphatidylcholine 17:0/17:0 (PC), phosphatidylethanolamine 17:0/17:0 (PE), lysophosphatidylcholine 12:0, (LPC) lysophosphatidylethanolamine 17:1 (LPE), triacylglycerol 17:0/17:0/17:0 (TAG) and sphingomyelin 18:1;2/12:0 (SM).

Samples were analyzed by direct infusion in a QExactive mass spectrometer (Thermo Scientific) in a single acquisition. Tandem mass-spectrometry (MS/MS) was triggered by an inclusion list encompassing corresponding MS mass ranges scanned in 1 Da increments. MS and MS/MS data were combined to monitor CE, DAG and TAG ions as ammonium adducts; PC, PC O-, as acetate adducts; and PE, PE O- and PI as deprotonated anions. MS only was used to monitor LPE as deprotonated anion; Cer, SM and LPC as acetate adducts and cholesterol as ammonium adduct.

Data post-processing and normalization were performed using an in-house developed data management system. Only lipid identifications with a signal-to-noise ratio >5 and a signal intensity 5-fold higher than in corresponding blank samples were considered for further analysis.

#### Targeted sphingolipid measurements

Ceramides (C16:0 cer, C18:0 cer, C18:1 cer, C20:0 cer, C22:0 cer, C24:0 cer and C24:1 cer), Dihydroceramides (C16:0 DHcer, C18:0 DHcer, C18:1 DHcer, C20:0 DHcer, C22:0 DHcer, C24:0 DHcer,C24:1 DHcer) and precursors (Sphingosine, Sphinganine, 1-Deoxysphinganine,1-Methyldeoxysphinganine, SB) were quantified in plasma by liquid chromatography tandem mass spectrometry (LC-MC/MC). In addition to samples, seven-point calibration curves and 3 levels of quality controls were made from pure standards in BSA 5%. Finally, reference plasma spiked with analytes at two different levels were prepared as additional QC samples.

After lipid chromatographic separation on a UPLC I-Class system (Waters), mass analysis was performed on an API 6500 system (Sciex) operating with an electrospray source in positive mode. General parameters were set as follows: curtain gas: N2 (35 PSI), Ion source gas 1: Air (50 PSI), Ion source gas 2: Air (50 PSI), ion source voltage: 5500 V, temperature: 300°C, collision gas: N2 (7). Scheduled multiple reaction monitoring (MRM) mode was used with a target scan time of 0.5s and an MRM detection window of 60s.

Data was acquired using Analyst 1.6.2 (Sciex) and data processing was performed with MultiQuant 3.0 (Sciex). Peak area of analyte and internal standard were determined by the MultiQuant 3.0 (Sciex) integration system. Analyte concentrations were determined using the internal standard method. The standard curves were generated from the peak area ratios of analyte/internal standard using linear regression analysis with 1/x2 weighting (except for C24 cer: quadratic regression analysis). Quantifications of analytes were accepted based on quality control samples. A tolerance of 25% and 30% was applied for accuracy and precision of QC samples and spiked plasma samples, respectively. All concentrations were reported in ng/mL.

#### Statistical analysis of shotgun lipidomics and targeted sphingolipid data

The statistical analyses of the shotgun lipidomics and targeted sphingolipid data sets were kept separate. Identical analysis steps were applied to the two data sets. Both sets had missing data values. Lipid species with ≥25% missing values across all available plasma samples were removed from the data set. This filtering resulted in 113 lipid species that were kept in the shotgun data set (523 were removed) and 14 in the targeted data set (4 were removed). For the lipids that remained in the data sets, missing values were imputed using a random forest approach, applying the function missForest from the R package missForest, with default parameters. In a next step, samples were filtered based on subject characteristics: individuals with bilirubin levels ≥100 μmol/l were removed before all analysis; moreover, individuals categorized as IGT with an HbA1c≥6% were excluded from the group comparisons in differential analysis, but they were retained in other analyses involving lipidomics data. In differential analysis, due to the limited number of available ND samples, the ND and the included IGT samples were combined into a single group for comparison with other sample groups, as described in the result section.

For differential analysis, linear models were applied, using the function lm from the R stats package. For each comparison between two sample groups, a linear model that included diabetes status as the main explanatory variable and age and sex as covariates was fitted to the data from the two groups. P values for diabetes status were adjusted across all included lipid species with the Benjamini-Hochberg method, separately for each comparison.

### Integrative analysis of transcriptomics and lipidomics

#### Multiblock modeling

Consensus Orthogonal Partial Least Squares (OPLS) model was computed with the MATLAB 9 environment with combinations of toolboxes and in-house functions that are available at https://gitlab.unige.ch/Julien.Boccard/consensusopls. Modified RV-coefficients were computed with the publicly available MATLAB m-file^50^. KOPLS-DA was assessed with routines implemented in the KOPLS open source package^51^. Consensus OPLS modeling was performed on shotgun lipidomics, targeted sphingolipids and transcriptomics data tables, which were all autoscaled prior to the analysis. The Consensus OPLS model distinguishes variation of data that is correlated to Y response and those which is orthogonal to Y response. This eases the biological interpretation of results and enables the link between variation of variables and variation of the outcome while removing information coming from other sources of variation.

The model resulted in 3 components, of which 1 predictive latent variable and 2 orthogonal latent variables. The quality of the model was assessed by *R*^2^ and *Q^2^* values, which define the portion of data variance explained by the model and the predictive ability of the model, respectively. The predictive component carried 11% of the total explained variance of global data (R^2^X) and explained 51.7% of variation of HbA1c (R^2^Y). This indicates that the model was able to explain a large part of variation of the response variable based on the different data matrices. The Q^2^ value was computed by a K-fold cross validation (K=7), which led to a goodness of prediction of Q^2^ = 0.26.

To ensure the validity of the model, a series of 1,000 permutation tests were carried out by mixing randomly the original Y response (HbA1c patient values). The true model Q2 value was clearly distinguished and statistically different from the random models distribution (*p*<0.001, mean=-0.1778, standard deviation (SD)=0.150, n=1,000). The variable relevance to explain the HbA1c variation was evaluated using the variable importance in projection (VIP) parameter, which reflects the importance of variables both with respect to the response and to the projection quality. The most relevant features were selected using a VIP threshold > 1.2.

### Proteomics

#### Sample Preparation

Pooled pancreatic islet cells with an approximate surface area of 80,000 μm^2^ were collected via Laser Capture Microdissection (LCM) onto adhesive cap tubes. Isolates were reconstituted in a 20 μl lysis buffer (PreOmics, Germany) and transferred into PCR tubes^52^. Samples were boiled at 95°C for 1min to denature proteins and reduce and alkylate cysteines without shaking in a thermocycler (Eppendorf GmbH) followed by sonication at maximum power (Bioruptor, Diagenode, Belgium) for 10 cycles of 30sec sonication and 30sec cooldown each. Sample liquid was briefly spun down and boiled again for 10min without shaking. 20μl of 100mM TrisHCl pH 8.5 (1:1 v/v) and 20ng Trypsin/LysC were added to each sample, followed by overnight digestion at 30°C without shaking. Next day, 40μl 99% Isopropanol 5% Trifluoroacetic acid (TFA) (1:1 v/v) was added to the solution and mixed by sonication. Samples were then subjected to stage-tip cleanup via styrenedivinylbenzene reversed-phase sulfonate (SDB-RPS). Sample liquid was loaded on one 14-gauge stage-tip plug. Peptides were cleaned up with 2×200μl 99% Isopropanol 5% TFA and 2×200μl 99% ddH2O 5% TFA in an in-house made Stage-tip centrifuge at 2,000xg, followed by elution in 40μl 80% Acetonitrile, 5% Ammonia and dried at 45°C in a SpeedVac centrifuge (Eppendorf, Concentrator plus) according to the ‘in-StageTip’ protocol (PreOmics, Germany). Peptides were resuspended in 0.1% TFA, 2% ACN, 97.9% ddH2O.

#### Liquid chromatography and mass spectrometry (LC-MS)

LC-MS was performed with an EASY nanoLC 1200 (Thermo Fisher Scientific) coupled online to a trapped ion mobility spectrometry quadrupole time-of-flight mass spectrometer (timsTOF Pro, Bruker Daltonik GmbH, Germany) via nano-electrospray ion source (Captive spray, Bruker Daltonik GmbH). Peptides were loaded on a 50cm in-house packed HPLC-column (75μm inner diameter packed with 1.9μm ReproSil-Pur C18-AQ silica beads, Dr. Maisch GmbH, Germany). Sample analytes were separated using a linear 120min gradient from 5-30% buffer B in 95min followed by an increase to 60% for 5min, and by a 5min wash at 95% buffer B at 300nl/min (Buffer A: 0.1% Formic Acid, 99.9% ddH2O; Buffer B: 0.1% Formic Acid, 80% CAN, 19.9% ddH2O). The column temperature was kept at 60°C by an in-house manufactured oven.

Mass spectrometry analysis was performed in a data-dependent PASEF mode with 1 MS1 survey TIMS-MS and 10 PASEF MS/MS scans per acquisition cycle. Ion accumulation and ramp time in the dual TIMS analyzer was set to 100ms each and we analyzed the ion mobility range from 1/K_0_ = 1.6 Vs cm^-2^ to 0.6 Vs cm^-2^. Precursor ions for MS/MS analysis were isolated with 2Th windows for m/z<700 and 3Th for m/z>700 in a total m/z range of 100-1,700 by synchronizing quadrupole switching events with the precursor elution profile from the TIMS device. The collision energy was lowered linearly as a function of increasing mobility starting from 59 eV at 1/K_0_=1.6 VS cm^-2^ to 20 eV at 1/K_0_=0.6 Vs cm^-2^. Singly charged precursor ions were excluded with a polygon filter (otof control, Bruker Daltonik GmbH). Precursors for MS/MS were picked at an intensity threshold of 2.500 a.u. and resequenced until reaching a ‘target value’ of 20,000 a.u taking into account a dynamic exclusion of 40sec elution^24^.

#### Proteomics raw file processing

Raw files were searched against the human Uniprot databases (UP000005640_9606.fa, UP000005640_9606_additional.fa) MaxQuant (Version 1.6.7), which extracts features from four-dimensional isotope patterns and associated MS/MS spectra^53^. False-discovery rates were controlled at 1% both on peptide spectral match (PSM) and protein level. Peptides with a minimum length of seven amino acids were considered for the search including N-terminal acetylation and methionine oxidation as variable modifications and cysteine carbamidomethylation as fixed modification, while limiting the maximum peptide mass to 4,600 Da. Enzyme specificity was set to trypsin cleaving c-terminal to arginine and lysine. A maximum of two missed cleavages were allowed. Maximum precursor and fragment ion mass tolerance were searched as default for TIMS-DDA data, while the main search peptide tolerance was set to 20ppm. The median absolute mass deviation for the data was 0.68ppm. Peptide identifications by MS/MS were transferred by matching four-dimensional isotope patterns between the runs with a 0.7-min retention-time match window and a 0.05 1/K_0_ ion mobility window^54^. Label-free quantification was performed with the MaxLFQ algorithm and a minimum ratio count of 1^55^.

#### Bioinformatic analysis

Bioinformatics analysis was performed in Perseus (version 1.6.7.0 and 1.5.5.0) and GraphPad Prism (version 8.2.1)^56^. Reverse database, contaminant, and only by site modification identifications were removed from the dataset. Data were grouped by analytical replicates and filtered to at least 70% data completeness in one group. Missing values were imputed from a normal distribution with a downshift of 1.8 and a width of 0.3 and data were log_2_-transformed. To represent the data reproducibility and variability, a principal component analysis was performed on the median data of analytical replicate measurements of each individual. Clinically classified T2D and ND individuals were tested for differences in their mean by a two-sided Student’s t-test with S0=0.1 and a Benjamini-Hochberg correction for multiple hypothesis testing at an FDR of 0.05 preserving grouping of each individuals analytical replicate measurements, and presented as volcano plot. We then normalized the data by row-wise z-scoring followed by hierarchical clustering using Euclidean as the distance parameter for column- and row-wise clustering. 1D gene ontology enrichments of clustered and systematically changed proteins were performed with regards to their cellular compartment and keywords assignment^28^. Log2 transformed LFQ data were used for the calculation of intensity shifts of the enriched keyword or cellular compartment term for each of the displayed clusters. Total protein copy number estimation of the median LFQ intensities for patients clinically classified as non-diabetic and diabetic were calculated using the Perseus plugin ‘Proteomic ruler’^27^. Median LFQ intensity values for all T2D and ND were calculated. We annotated protein groups for the leading protein ID with the human Uniprot fasta file (UP000005640_9606.fa) and estimated the protein copy number with the following settings: Averaging mode. ‘All columns separately’, Molecular masses: ‘Average molecular mass’, Detectability correction: ‘Number of theoretical peptides’, Scaling mode: ‘Histone proteomic ruler’, Ploidy: ‘2’, Total cellular protein concentration: ‘200g/l’. Proteins were annotated with regards to their cellular compartment by gene ontology. We calculated the median protein copy number for the samples from T2D and ND PPP separately and multiplied it by its protein mass. To calculate the subcellular protein mass contribution, we calculated the protein mass proportion for the GOCC terms ‘Nucleus’, ‘Mitochondrion’, ‘Cytoskeleton’, ‘Golgi apparatus’, and ‘Endoplasmic reticulum’. For calculating the organellar change between T2D and ND PPP, protein mass contributions of each organelle were normalized by its respective ‘Nuclear part’ contribution. Chromosomal annotation of significantly changed proteins between T2D and ND PPP was identified via Ensembl ID.

### Antibody validation

Rabbit polyclonal anti-ALDOB antibody (Proteintech, Cat.No. 18065-1-AP) was tested for specificity by western blotting of protein extracts of *ALDOB^-/-^* MIN6 cells generated with a CRISPR/Cas9 system, as described^52^. The knock-out of *ALDOB* was verified by Sanger sequencing of the target locus.

### Isolated mouse islet and cell line experiments

Mouse (C57Bl6, db/db and db/+ mice, 3 animals/strain, age 13 weeks) islets were cultured for 1 day post isolation. Islet beta MIN6s4 and alpha αTC1-clone 6 cell lines were harvested for RNA extraction using Qiagen RNeasy Mini Kit according to the manufacturer’s instructions. After quality control, RNA samples were sequenced using the Illumina HiSeq 2000 platform and processed as previously described^45,57,58^.

### Immunofluorescence microscopy

Immunofluorescence staining was done on formalin-fixed paraffin embedded 5μm thick sections of human pancreatic tissue. Acetylated histone H3 and H4 were detected in separate sections using rabbit polyclonal antibodies (Merck Millipore Cat.No. 06-598 and 06-599, respectively). A mouse monoclonal anti-insulin antibody (Thermo Fisher Scientific Cat.No. 53-9769-82) was used for co-staining, to identify the beta cell areas. Images were acquired using a Nikon C2+ confocal microscope with a 60x oil immersion objective, with acquisition parameters normalized to a negative control sample.

---

**Figure S1:**
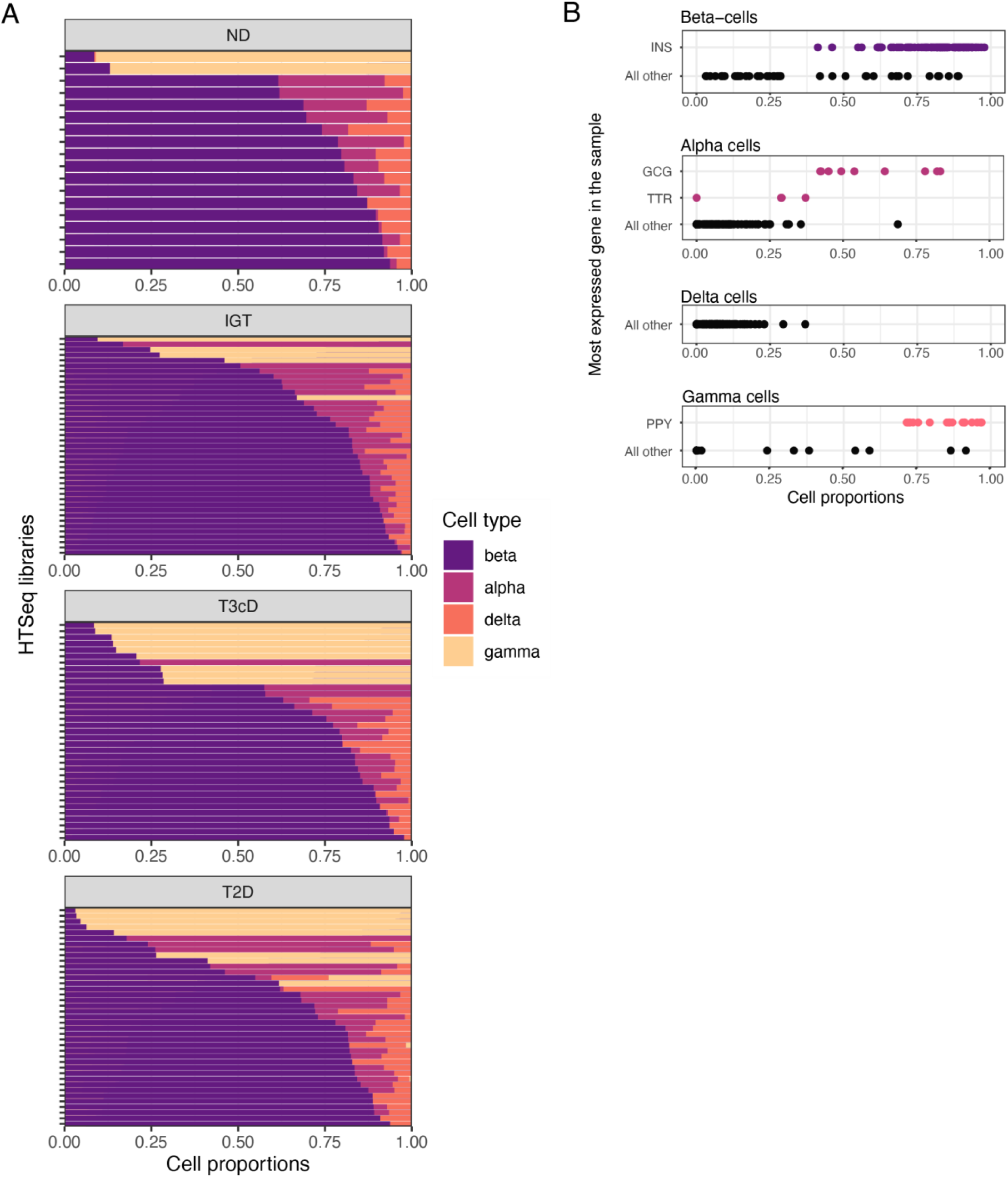
Deconvolution of cell types based on RNA-Seq data. A) Cell-type proportions by sample, as estimated with DeconRNASeq, panels faceted according to diabetes status. B) Sample distribution across each cell type proportion. Highlighted are samples presenting a cell type specific gene being the most expressed. Marker genes were *GCG* and *TTR* for alpha cells, *INS* for beta cells, *SST* for delta cells, and *PPY* for gamma cells.

**Figure S2:**
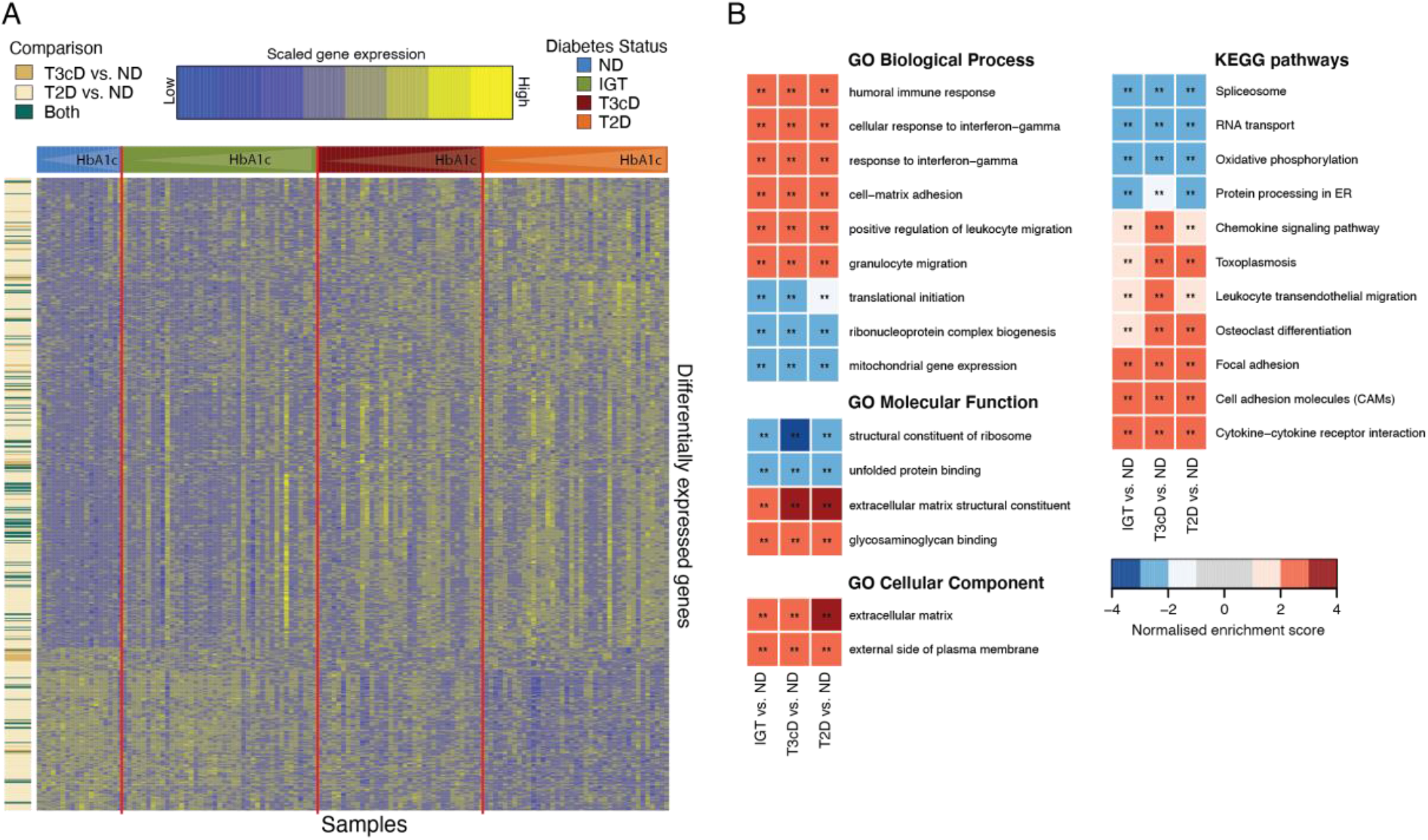
Differential gene expression analysis between glycemic groups in the entire cohort. A-B) Gene expression profile (A) and GSEA analysis (B) of DE genes between IGT, T3cD or T2D and ND PPP. Results are similar to those shown in Fig. 2BC, but obtained from the entire cohort of 133 PPP.

**Figure S3:**
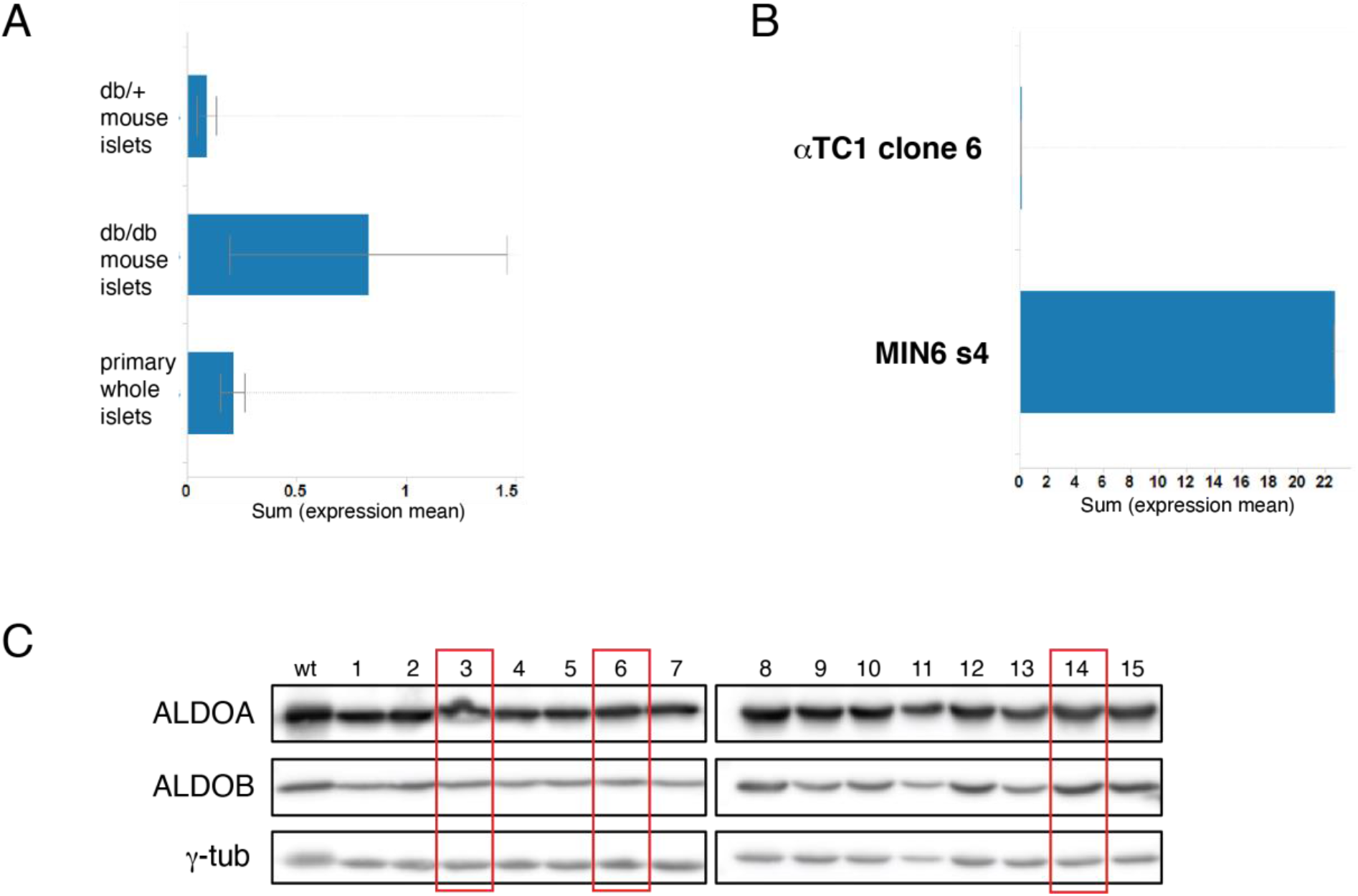
A-B) *ALDOB* expression (RNAseq, Illumina) in (A) islets from 13-week-old *db/db, db/+* mice and *C57Bl6* mice (3 animals/strain) or (B) mouse αTC1 clone 6 alpha and Min6s4 beta cell lines (n=4/cell line). C) Western blot of MIN6 single cell-derived clones with antibodies against ALDOB and ALDOA. Framed lanes mark *ALDOB* knockout clones as verified by site-specific sequencing.

**Figure S4:**
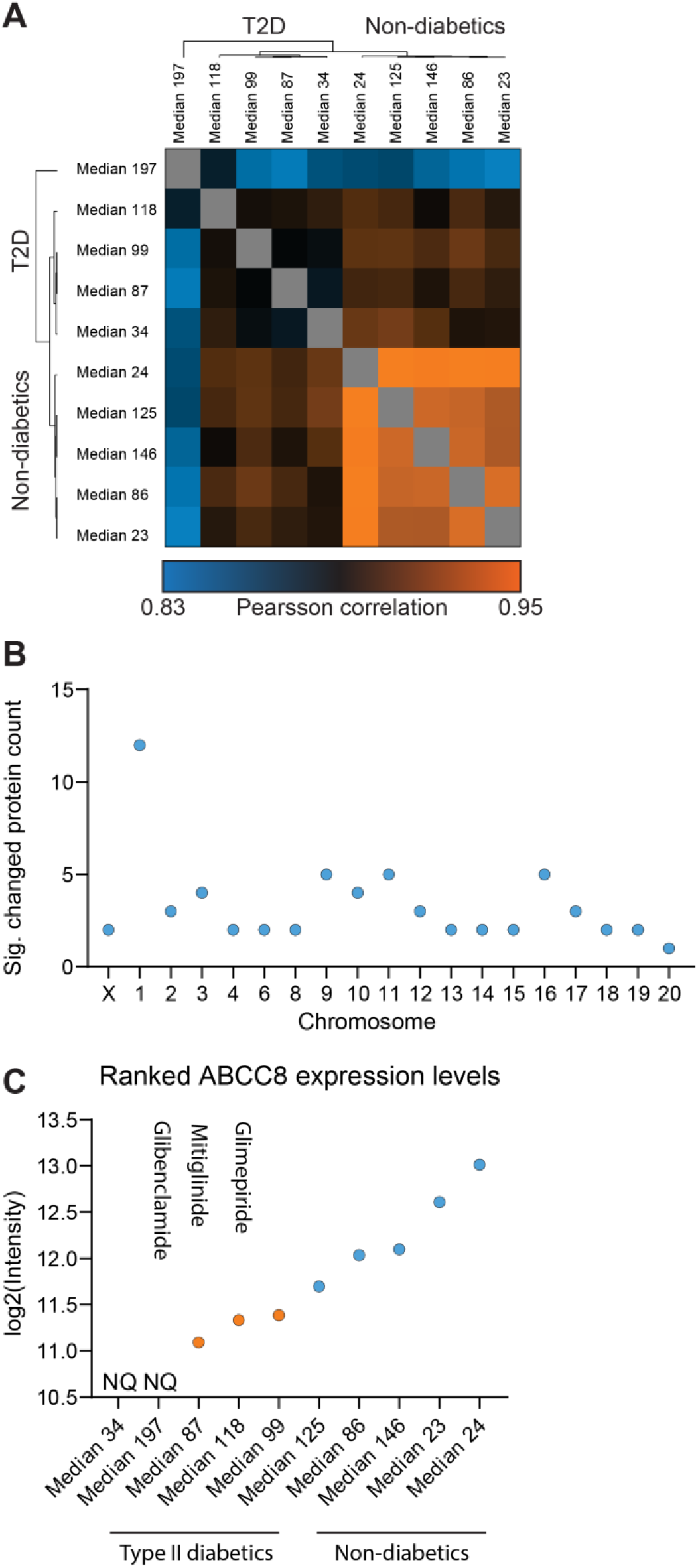
Hierarchical clustering of protein expression correlations in all biological replicates highlighting the technical and biological reproducibility of our proteome data set (A). Distribution of differentially expressed proteins between T2D and ND across chromosomes (B). Ranked ABCC8 protein expression levels across T2D and ND subjects. T2D are highlighted in orange, ND are highlighted in blue. Patient 118 was treated with Glimepiride; Patient 87 was treated with Mitiglinide; Patient 197 was treated with Glibenclamide (C).

## Supplementary Table Legends

**Table S1:** Clinical characteristics of the complete cohort of PPP for LCM islet RNA sequencing. Except for absolute frequencies, all values are mean ± standard deviation. The statistical testing was performed with two-sided t-Test comparisons with ND (\**p*<0.05, \*\**p*<0.01, \*\*\**p*<0.001).

**Table S2:** Differentially expressed (DE) islet genes between T3cD and T2D PPP compared with ND PPP in the entire cohort. No DE islet genes were identified when comparing IGT and ND PPP. Genes were considered differentially expressed when the adjusted p value was ≤ 0.05 and the fold change > 1.5.

**Table S3:** Clinical characteristics of the LCM islet RNA sequencing cohort with *INS* as the highest expressed gene. Except for absolute frequencies, all values are mean ± standard deviation. Statistical testing was performed with two-sided t-Test comparisons with ND (\**p*<0.05, \*\**p*<0.01, \*\*\**p*<0.001).

**Table S4:** Differentially expressed (DE) islet genes between IGT, T3cD or T2D PPP compared with ND PPP in the “restricted” cohort. Genes were considered differentially expressed when the adjusted *p* value was ≤ 0.05 and the fold change > 1.5.

**Table S5:** Complete results of KEGG pathways and GO term gene set enrichment analyses of differentially expressed genes between glycemic groups in the “restricted” PPP cohort.

**Table S6:** Complete results of KEGG pathways and GO term gene set enrichment analyses of differentially expressed genes between glycemic groups in the entire PPP cohort.

**Table S7:** Significance of co-expressed gene modules.

**Table S8:** Clinical characteristics of the PPP cohort for proteomic analyses.

**Table S9:** Clinical characteristics of the PPP cohort for shotgun lipidomic analyses.

**Table S10:** Shotgun lipidomics. Lipid classes and number of species per class included in the data analysis.

**Table S11:** Clinical characteristics of the PPP cohort for sphingolipid analyses.

**Table S12:** Targeted lipidomics. Names of ceramide and sphingolipid classes included in the data analysis.

**Table S13:** Result lists of differential analysis, sorted by *p* value, in plasma shotgun lipidomic data (first two Excel sheets) and in targeted sphingolipid data (last two Excel sheets), from comparisons of T2D vs. ND and T3cD vs. ND PPP, with ND as defined in lipidomics result section. All lipid species that were included in the analysis are shown. Mean lipid concentrations were considered significantly different between groups when the adjusted *p* value was ≤ 0.05.

**Table S14:** Consensus OPLS predictive scores and loadings.

**Table S15:** Complete results of KEGG pathways over representation analyses of selected co-expressed gene modules.

