## Supplementary Tables for "Multi-omics profiling of living human pancreatic islet donors reveals heterogeneous beta cell trajectories toward type 2 diabetes": suppl_table_S1_fig1_cohortComplete.docx

t

|  | Gender (M/F) | Age  (years) | BMI | Fasting glucose (mmol/l) | HbA1c (%) | Glycaemia at 2h OGTT (mmol/l) | Underlying pancreatic disease | | |
| --- | --- | --- | --- | --- | --- | --- | --- | --- | --- |
|  |  |  |  |  |  |  | Chronic pancreatitis | Benign tumour | Malignant tumour |
| ND | 10/8 | 56.72±10.89 | 22.9±2.88 | 4.93±0.32 | 5.25±0.3 | 5.87±1.2 | 4  (22.22%) | 5 (27.78%) | 9  (50%) |
| IGT | 18/23 | 65.92±11.54** | 26.34±3.68*** | 5.6±0.59*** | 5.75±0.42*** | 8.27±1.7*** | 6  (14.63%) | 10  (24.39%) | 25  (60.97%) |
| T3cD | 18/17 | 67.45±9.55** | 26.03±4.36** | 6.55±1.39*** | 6.29±0.95*** | 11.82±2.54*** | 6  (17.14%) | 3  (8.57%) | 26  (74.29%) |
| T2D | 23/16 | 66.87±12.06** | 25.99±5.22** | 7.87±2.6*** | 7.41±1.29*** | NA | 6  (15.38%) | 6  (15.38%) | 27  (69.23%) |
