## Supplementary Tables for "Multi-omics profiling of living human pancreatic islet donors reveals heterogeneous beta cell trajectories toward type 2 diabetes": suppl_table_S3_fig1_INS_cohort.docx

|  | Gender (M/F) | Age  (years) | BMI | Fasting glucose (mmol/l) | HbA1c (%) | Glycaemia at 2h OGTT (mmol/l) | Underlying pancreatic disease | | |
| --- | --- | --- | --- | --- | --- | --- | --- | --- | --- |
|  |  |  |  |  |  |  | Chronic pancreatitis | Benign tumour | Malignant tumour |
| ND | 8/7 | 55.8±11.71 | 22.83±3.15 | 4.9±0.32 | 5.2±0.3 | 5.73±1.22 | 4 | 4 | 7 |
| IGT | 14/18 | 64.9±12.34* | 26.58±3.79** | 5.46±0.58*** | 5.75±0.39*** | 8.22±1.45*** | 6 | 7 | 19 |
| T3cD | 11/10 | 68.42±9.84** | 26.22±4.31* | 6.8±1.66*** | 6.62±0.91*** | 11.8±3.07*** | 3 | 0 | 18 |
| T2D | 12/12 | 67.04±12.7** | 26.58±4.8** | 8.47±2.48*** | 7.48±1.2*** | NA | 4 | 3 | 17 |
