## Supplementary Tables for "Multi-omics profiling of living human pancreatic islet donors reveals heterogeneous beta cell trajectories toward type 2 diabetes": suppl_table_S8_fig4_cohort proteomic.docx

|  | Gender (M/F) | Age  (years) | BMI | Fasting glucose (mmol/l) | HbA1c (%) | Glycaemia at 2h OGTT (mmol/l) | Underlying pancreatic disease | | |
| --- | --- | --- | --- | --- | --- | --- | --- | --- | --- |
|  |  |  |  |  |  |  | Chronic pancreatitis | Benign tumour | Malignant tumour |
| ND | 2/3 | 54.8±18.15 | 26.79±4.38 | 4.93±0.2 | 4.77±0.51 | 6.3±0.85 | 2 | 1 | 2 |
| T2D | 2/3 | 59.6±9.18 | 26.25±5.23 | 10.62±2.14 | 9±0.89 | NA | 1 | 1 | 3 |
