## Supplementary Tables for "Multi-omics profiling of living human pancreatic islet donors reveals heterogeneous beta cell trajectories toward type 2 diabetes": suppl_table_S9_fig5_cohort shotgun lipidomics.docx

|  | Gender (M/F) | Age  (years) | BMI | Fasting glucose (mmol/l) | HbA1c (%) | Glycaemia at 2h OGTT (mmol/l) | Underlying pancreatic disease | | |
| --- | --- | --- | --- | --- | --- | --- | --- | --- | --- |
|  |  |  |  |  |  |  | Chronic pancreatitis | Benign tumour | Malignant tumour |
| ND | 2/2 | 61.25±2.63 | 22.27±2.05 | 5.05±.03 | 5.47±0.12 | 6.41±0.86 | 0 | 1 | 3 |
| IGT | 9/12 | 71.38±7.91 | 27.13±4.12 | 5.77±0.66 | 5.81±0.44 | 8.61±1.47 | 1 | 5 | 15 |
| T3D | 5/8 | 70±9.74 | 24.11±4.74 | 8.55±7.86 | 6.5±1.75 | 13.06±3.17 | 2 | 2 | 9 |
| T2D | 6/11 | 66.88±12.62 | 25.67±5.8 | 8.21±2.17 | 7.4±1.16 | NA | 3 | 6 | 8 |

**Supplementary table S8** Clinical characteristics of the cohort for lipidomic analyses.
