## Supplementary Tables for "Multi-omics profiling of living human pancreatic islet donors reveals heterogeneous beta cell trajectories toward type 2 diabetes": suppl_table_S10_fig5 shotgun_lipidomics_classes.docx

| **Lipid Class** | **Number of species**  **in data analysis** |
| --- | --- |
| Ceramides (Cer) | 4 |
| Diacylglycerols (DAG) | 1 |
| Lysophosphatidylcholines (LPC) | 6 |
| Lysophosphatidylethanolamines (LPE) | 6 |
| Phosphatidylcholines (PC) | 6 |
| Ether-linked Phosphatidylcholines (PC O-) | 30 |
| Phosphatidylethanolamines (PE) | 6 |
| Ether-linked Phosphatidylethanolamines (PE O-) | 10 |
| Phosphatidylinositols (PI) | 5 |
| Sphingomyelins (SM) | 10 |
| Triacylglyerols (TAG) | 29 |
