## Supplementary Tables for "Multi-omics profiling of living human pancreatic islet donors reveals heterogeneous beta cell trajectories toward type 2 diabetes": suppl_table_S11_fig5 cohort targeted sphingolipids.docx

|  | Gender (M/F) | Age  (years) | BMI | Fasting glucose (mmol/l) | HbA1c (%) | Glycaemia at 2h OGTT (mmol/l) | Underlying pancreatic disease | | |
| --- | --- | --- | --- | --- | --- | --- | --- | --- | --- |
|  |  |  |  |  |  |  | Chronic pancreatitis | Benign tumour | Malignant tumour |
| ND | 6/5 | 57.36±10.21 | 23.19±2.81 | 4.94±0.28 | 5.3±0.25 | 6.14±0.84 | 3 | 3 | 5 |
| IGT | 13/19 | 66.81±11.85 | 26.44±3.66 | 5.67±0.63 | 5.72±0.44 | 8.12±1.82 | 3 | 9 | 20 |
| T3D | 14/12 | 67.88±9.8 | 25±4.22 | 7.74±5.54 | 6.51±1.43 | 12.38±2.53 | 5 | 2 | 19 |
| T2D | 15/17 | 67.09±12.9 | 26.1±5.79 | 8.09±2.52 | 7.13±1.19 | NA | 5 | 6 | 21 |
