## Supplementary Tables for "Multi-omics profiling of living human pancreatic islet donors reveals heterogeneous beta cell trajectories toward type 2 diabetes": suppl_table_S12_fig5 targeted sphingolipid classes.docx

| **Ceramide/Sphingolipid Classes** | **Lipid Species** | **Alternative name** |
| --- | --- | --- |
| Ceramides | C16 Cer | Cer(d18:1/16:0) |
|  | C18 Cer | Cer(d18:1/18:0) |
|  | C20 Cer | Cer(d18:1/20:0) |
|  | C22 Cer | Cer(d18:1/22:0) |
|  | C24 Cer | Cer(d18:1/24:0) |
|  | C24:1 Cer | Cer(d18:1/24:1) |
| Dihydroceramides | C16 DH Cer | Cer(d18:0/16:0) |
|  | C18 DH Cer | Cer(d18:0/18:0) |
|  | C20 DH Cer | Cer(d18:0/20:0) |
|  | C22 DH Cer | Cer(d18:0/22:0) |
|  | C24 DH Cer | Cer(d18:0/24:0) |
|  | C24:1 DH Cer | Cer(d18:0/24:1) |
| Sphingoid bases | Sphinganine | Sphinganine |
|  | Sphingosine | Sphing-4-enine |
